# Mitotic R-loops direct Aurora B kinase to maintain centromeric cohesion

**DOI:** 10.1101/2021.01.14.426738

**Authors:** Erin C. Moran, Limin Liu, Ewelina Zasadzinska, Courtney A. Kestner, Ali Sarkeshik, Henry DeHoyos, John R. Yates, Daniel Foltz, P. Todd Stukenberg

## Abstract

Recent work has shown that R-loops exist at mitotic centromeres, but the function of these R-loops is not well understood. Here, we report that mitotic R-loops arise in distinct locations from those formed during interphase. They accumulate on chromosome arms in prophase, where they are quickly resolved and continue to be produced at repetitive sequences including centromeres during a mitotic stall. Aurora B kinase activity is required to resolve R-loops during prophase and R-loops promote the localization of the Chromosome Passenger Complex (CPC) to the inner centromere. CPC purified from mitotic chromosomes interacts with thirty-two proteins involved with R-loop biology. One of these, the RNA regulator RBMX, controls Aurora B localization and activity in vivo. Perturbations in R-loop homeostasis or RBMX cause defects in the maintenance of centromeric cohesion due to the mislocalization of the CPC. We conclude that R-loops are generated by mitotic processes in repetitive DNA sequences, they play important roles in mitotic fidelity, and we have identified a set of mitotic R-loop regulators including the CPC and RBMX that will enable future studies of mitotic R-loops.

## Introduction

Mitosis is characterized by a reorganization of chromatin structure as most transcription is silenced and terminated (Gottesfeld and Forbes, 1997; Jiang et al., 2004; Taylor, 1960), the higher order structure of chromatin such as topologically associating domains (TADs) are removed (Naumova et al., 2013), and chromatin is condensed. R-loops are three stranded structures in which a strand of RNA is annealed to a denatured double-stranded DNA. R-loops have distinct roles in interphase cells such as transcriptional initiation and termination (Ginno et al., 2012, 2013; Skourti-Stathaki et al., 2014), chromatin compaction (Castellano-Pozo et al., 2013a; Nakama et al., 2012), and both as a source of DNA damage (Bhatia et al., 2014; Costantino and Koshland, 2018; Gan et al., 2011; Wahba et al., 2011) and DNA damage repair (Ohle et al., 2016; Yasuhara et al., 2018). In mitotically arrested cells, R-loops localize to centromeres, are associated with a centromeric DNA damage response (Kabeche et al., 2018) and form the basis of centromeric chromatin loops in maize cells (Liu et al., 2020). However, the function of R-loops at the centromere is poorly understood, and there are major unanswered questions including: whether R-loops arise during an unperturbed mitosis, if they arise only in centromeric sequences, if R-loops are resolved as mitosis progresses, and what is the consequence of not generating mitotic R-loops to mitotic cells.

Aurora B kinase-dependent H3S10 phosphorylation increases in *S. cerevisiae* strains that accumulate R-loops (Castellano-Pozo et al., 2013a), however it is not clear if Aurora kinases play an active role in regulation of R-loops. It was recently reported that mitotic R-loops are required to activate Aurora B kinase, a major regulator of mitotic events, and this signal is downstream of ATR and CHK1 signaling in the DNA damage response (Kabeche et al., 2018). Aurora B kinase is directly phosphorylated by CHK1 to regulate kinase activity (Petsalaki et al., 2011). Aurora B kinase regulates multiple important steps in mitosis, such as the spindle assembly checkpoint (Biggins and Murray, 2001; Hauf et al., 2003; Kallio et al., 2002; Sacristan and Kops, 2015; Santaguida et al., 2011; Stukenberg and Burke, 2015), sister chromatid cohesion (Dai et al., 2006; Resnick et al., 2006; Tanno et al., 2010), and kinetochore-microtubule attachment regulation (Cimini et al., 2006; Knowlton et al., 2006; Liu et al., 2009; Meppelink et al., 2015; Salimian et al., 2011; Welburn et al., 2010). Aurora B kinase associates with three other proteins to form the Chromosome Passenger Complex (CPC). The CPC has a very dynamic localization during prophase; it is initially localized throughout condensing chromosomes, transitions to the axis between sister chromatids, then to the inner centromere (Carmena et al., 2012; Hindriksen et al., 2017; Hirota et al., 2005; Jeyaprakash et al., 2007; Klein et al., 2006; Nozawa et al., 2010). One function of Aurora B on chromosome arms is to remove most interphase cohesin, which is likely the pool at the base of TADs (Losada et al., 2005; Naumova et al., 2013). Paradoxically, sister chromatid cohesion is maintained at inner centromeres, where Aurora B is highest during late prophase until anaphase onset. This is due to a Sgo1-dependent mechanism, whereby Sgo1 recruits the B56-PP2A phosphatase complex in order to maintain cohesion (Kang et al., 2011; Kitajima et al., 2006; Meppelink et al., 2015; Tang et al., 2006). Centromeric CPC is required to maintain sister chromatid cohesion at the centromere(Dai et al., 2006; Resnick et al., 2006), but the mechanism of this activity is less clear.

There is growing evidence that transcripts are made from centromeres in multiple species, but the functions of these transcripts are poorly understood. The human centromere is made up of alpha satellite DNA, forming <u>H</u>igher <u>O</u>rder <u>R</u>epeat (HOR) structures (Willard, 1985). HORs are flanked by pericentric heterochromatin, which is formed by alpha satellite monomers and other satellite sequences such as beta and gamma satellites, as well as Human Satellite I and II sequences. The alpha satellite HOR sequences are transcribed in G2/M cells (Hall et al., 2014; Ideue et al., 2014; Liu et al., 2015). Active RNA Polymerase II has been detected at human centromeres in multiple circumstances (Chan et al., 2012; Liu et al., 2015; McNulty et al., 2017); this transcription is well conserved across species. These RNAs are known to be spliced in some organisms (Grenfell et al., 2016; Liu et al., 2020), and in *Xenopus laevis* these spliced transcripts are known to interact with the CPC, which is required to localize the CPC (Blower, 2016; Grenfell et al., 2016; Jambhekar et al., 2014). Recent efforts to identify RNAs associated with chromatin has also shown that RNA made from human centromeres associate with the chromatin in *cis* (McNulty et al., 2017) and have long half-lives (Hall et al., 2014; McNulty et al., 2017) suggesting they are stabilized in some way. Multiple components of the human spliceosome have now been found to be important for cohesion (Huen et al., 2010; Karamysheva et al., 2015; van der Lelij et al., 2014; Nishimura et al., 2019; Sundaramoorthy et al., 2014), but the mechanism of how these proteins regulate cohesion is unclear. Recent efforts to identify proteins associated with R-loops (Cristini et al., 2018) have also identified many splicing proteins, suggesting that the mechanism of activity for these proteins involves R-loop homeostasis during mitosis.

We present evidence that R-loops accumulate on chromosomes during prophase, that they are resolved during mitosis and this resolution requires the CPC. In addition, R-loops are required to promote CPC localization to the inner centromere. We purified the CPC from mitotic chromosomes and identified 32 proteins that interact with R-loops, including the cohesion regulator RBMX. We show that R-loops and RBMX are required to localize the CPC to inner centromeres and this promotes localization of Sgo1 to protect centromeric cohesion. This work provides a function for mitotic R-loops that is distinct from interphase roles and provides a new function for the pool of chromatin-based Aurora B kinase in resolving R-loops in mitosis.

## Results

### R-loops are found at repetitive regions in mitotic cells

We investigated the genomic loci on mitotic chromosomes that contain R-loops and further tested whether there is a population of R-loops that are regulated by Aurora B kinase (Castellano-Pozo et al., 2013). Specifically, we performed DNA-RNA IP using S9.6 monoclonal antibody, which recognizes RNA-DNA hybrids (Boguslawski et al., 1986), followed by high throughput sequencing (DRIP-seq) on cells stalled in mitosis for 24 hours with colcemid and then treated with either DMSO, or two different Aurora B inhibitors AZD1152/Barasertib (AZD) or ZM447439 (ZM) for the last hour. Each sample had a paired RNaseH1 treated sample to show specificity of the IP. The DNA sequence reads were mapped to the human genome (hg38) and peaks were called by MACS2 (Feng et al., 2012; Zhang et al., 2008). We validated that the peaks were at R-loops by comparing an average line profile of the peaks in each treatment to the average line profile in the RNaseH treated control IPs. An example peak profile is shown in Figure 1A, where the called compiled peaks (~90,000 peaks per treatment) were approximately ten-fold enriched relative to their respective RNaseH1-treated sample.

**Figure 1.**
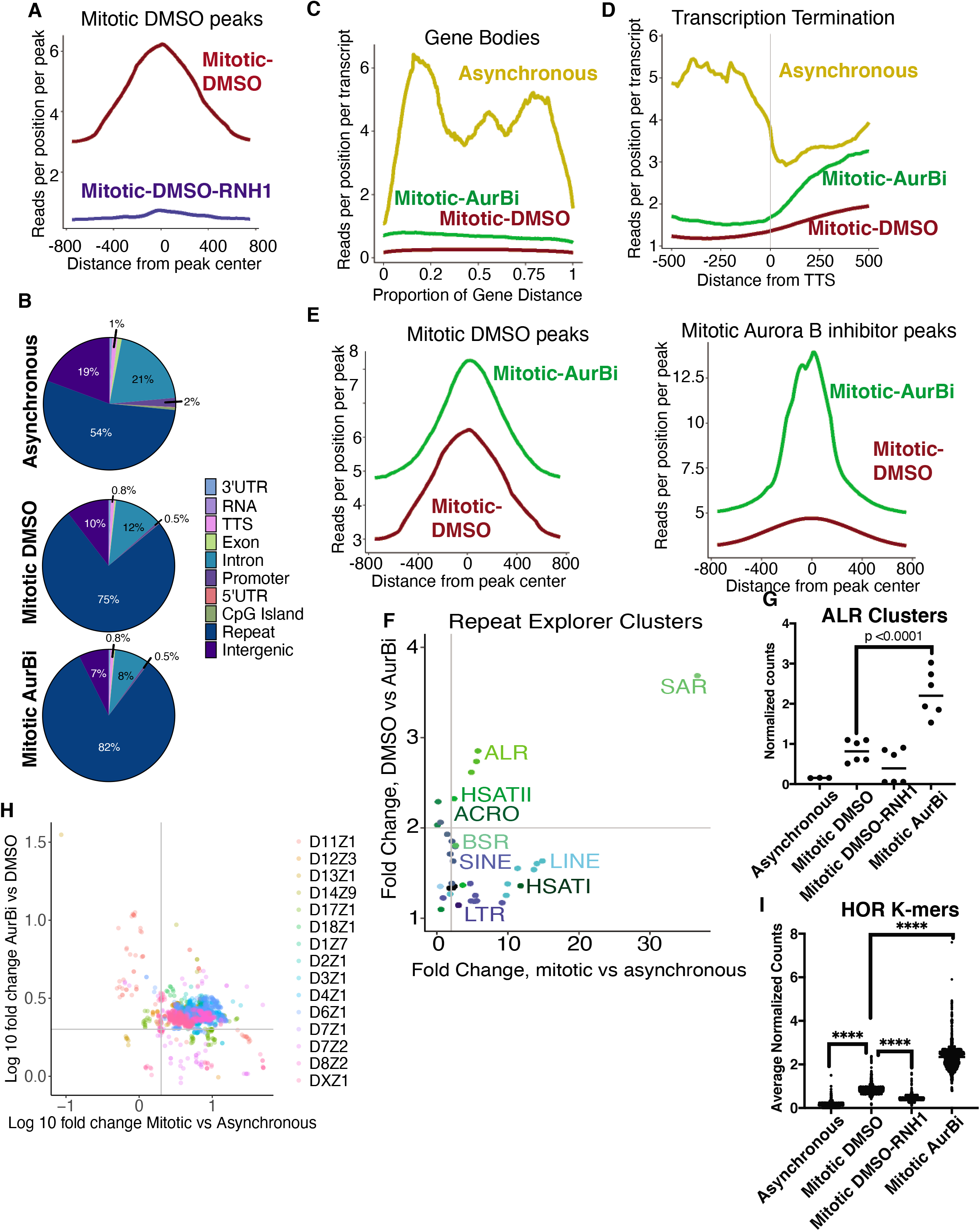
Aurora B is responsible for removing R-loops from Centromeric Satellite Repeats. DRIP-seq analysis of mitotic DLD1 cells, stalled in mitosis and treated with vehicle (DMSO), 500 nM AZD-1152, or 2 μM ZM-447439 for the last hour of the arrest, compared with asynchronous DRIP-seq from HEK293 cells published in (Nadel et al., 2015). A. Example track of peaks called in vehicle sequence samples, compared with paired RNaseH1-treated samples; replicate data compiled in each. Two DRIP-seq samples per track, measured across 96,139 loci. B. Enrichment of reads at peak loci called in vehicle samples (left, 96,139 loci) or Aurora B inhibited samples (right, 89,803 loci) with replicate data compiled. C. Genomic annotation of all peaks called by the same pipeline in asynchronous cells (28,573 loci), vehicle treatment (96,139 loci), and Aurora B inhibited (89,803 loci). D-E. Enrichment of reads across gene bodies, scaled to be a proportion of the gene length in D, and as measured near transcriptional termination sites (TTS) in E. Genes were identified via GENCODE v32. Yellow, Asynchronous cells, Red, Mitotic DMSO treated cells, Green, Mitotic Aurora B inhibited cells. F-G. Analysis of enrichment of reads from Repeat Explorer clusters, after filtering those clusters which had insufficient read counts and those with less than 2-fold enrichment relative to RNaseH1 treated samples. 100 clusters analyzed, 38 remained after filtering. Grey lines in F represent 2-fold enrichment. G. Analysis of clusters identified as ALR, 3 clusters measured over 2 DRIP-seq samples; plot shows individual measurements and mean values. H-I. Relative enrichment analysis of higher order repeat specific K-mers derived from Miga, 2017; 2119 measured in all. H. Individual k-mer enrichment log 10-fold changes in mitotic cells vs asynchronous cells on the x axis, Aurora B inhibited mitotic cells vs DMSO mitotic cells on the y axis, individual HORs in different colors. Grey lines, 2-fold enrichment lines. I. Distribution of counts of the total k-mer array, points show individual measurements and line represents the mean values. ****, p<0.0001, calculated by one-way ANOVA.

We first asked if R-loops are in the same locations as in interphase cells or whether new R-loops are formed during mitosis. We compared the called peaks of our mitotic DRIP-seq samples to publicly available data from asynchronous cells that were prepared using a similar IP protocol, sequenced to a similar depth and in cells with a similar karyotype as the cells used in our experiment (Nadel et al., 2015). Significantly more R-loops were found on repeat elements in mitotic cells compared to interphase cells. In addition, there was a decrease in peaks in promoters and genes in mitotic cells (Figure 1B), which included a drastic loss of R-loop accumulation across gene bodies and transcriptional termination sites in the mitotic samples (Figure 1C, D). The repression of DRIP signal within mitotic cells is consistent with the silencing of transcription during mitosis (Parsons and Spencer, 1997; Prescott and Bender, 1962; Taylor, 1960). These data suggest that mitotic R-loops are distinct from interphase R-loops, which is a consequence of the general termination of transcription that happens as cells enter mitosis and an accumulation of new R-loops at repetitive regions in the genome.

We next determined if peaks were regulated by Aurora B. Peaks that existed in mitotic control samples were more enriched after treatment with Aurora inhibitors (Figure 1E, left) and new peaks also appeared (Figure 1E, right). A larger percentage of the R-loops were found at repetitive elements after the addition of Aurora inhibitors for 1 hour (Figure 1B). There was not a dramatic change of R-loops in gene bodies or transcriptional start sites after Aurora B inhibition (Figure 1C, D) however, there was an increase in the number of R-loops at transcription termination sites. This is consistent with the recently identified role of Aurora B as a driver of the mitotic termination of transcription by its ability to remove cohesin from chromatin (Perea-Resa et al., 2020).

We focused on the enrichment of R-loops at repetitive elements in mitotic cells, since there was both a drastic increase in peaks called within repetitive elements in the mitotic samples and these further increased upon treatment with Aurora B inhibitors (Figure 1E). We devised a pipeline to identify the repetitive elements that were enriched in our DRIP-seq samples. The input samples were run through a De Bruijn graph algorithm to build a de novo database of repeat elements from DLD1 cells (Novák et al., 2010, 2013). We then aligned our DRIP-seq samples to this database and calculated enrichment values for each repetitive element. The repetitive elements were defined using the Dfam database (Hubley et al., 2016), and classified as subtype Transposable elements (Figure 1F, blue scale), subtype Satellite (green scale) or neither (black). R-loops accumulated (greater than 2-fold) at alpha-satellite repeats (ALR), scaffold attachment repeats (SAR), and Human satellite II repeats (HSATII) in mitosis and these further accumulated after Aurora B inhibition (Figure 1F). Our pipeline identified three distinct alpha-satellite clusters that represent slightly different sequences. All three were enriched in mitosis and further enriched after Aurora B inhibition with both AZD and ZM (treated as duplicates, averaged in Figure 1F, and shown in 1G, p<0.001). We determined the relative frequency of a set of published HOR-specific 24-base k-mers (Miga, 2017) in asynchronous, DMSO treated mitotic and Aurora B inhibited mitotic samples to test this enrichment in a complimentary manner. Almost all of the alpha-satellite k-mers were enriched greater than 2-fold in mitosis and these were further enriched after treatment with Aurora B inhibitors (Figure 1H, I, p<0.0001 for each pair in I). This enrichment is not apparent in samples treated with RNaseH, demonstrating specificity of the alpha-satellite enrichment. We conclude that R-loops accumulate in repetitive elements including alpha-satellite sequences during mitosis, and that Aurora B activity is required to deplete these R-loops.

### Dynamics of mitotic R-loops

We localized R-loops by immunofluorescence using the S9.6 monoclonal antibody on randomly cycling RPE1-T-REx cells to measure the dynamics of mitotic R-loops in a non-arrested cell population (Figure 2A). Cells were co-stained using Anti-Centromere Antibody (ACA) and the DNA-specific dye 4’,6-diamidino-2-phenylindole (DAPI). We quantified the R-loop signal intensity at centromeres and chromatin, using a 3D mask containing ACA signal to specify centromeres and a 3D mask created by DAPI staining as a chromatin signal. We then extrapolated the chromosome arm signal by subtraction of the centromere area from the chromatin mask. R-loop signals at both locations are highest in prophase cells, followed by a gradual decline in R-loop signal through the course of mitosis. The differences in centromeric R-loop intensities were significant in each of the distinct stages of mitosis, and the levels of centromeric R-loops had returned to interphase levels by anaphase (Figure 2B and Supplemental figure 1). Chromosome arm R-loops were also highest in prophase, but declined rapidly, to the point where they were statistically indistinguishable from interphase by metaphase (Figure 2C and Supplemental figure 1). This suggests that R-loops form during chromosome condensation across chromosomes but are restricted to centromeres after prophase. Centromeric R-loops persist longer but are resolved by anaphase onset.

**Figure 2.**
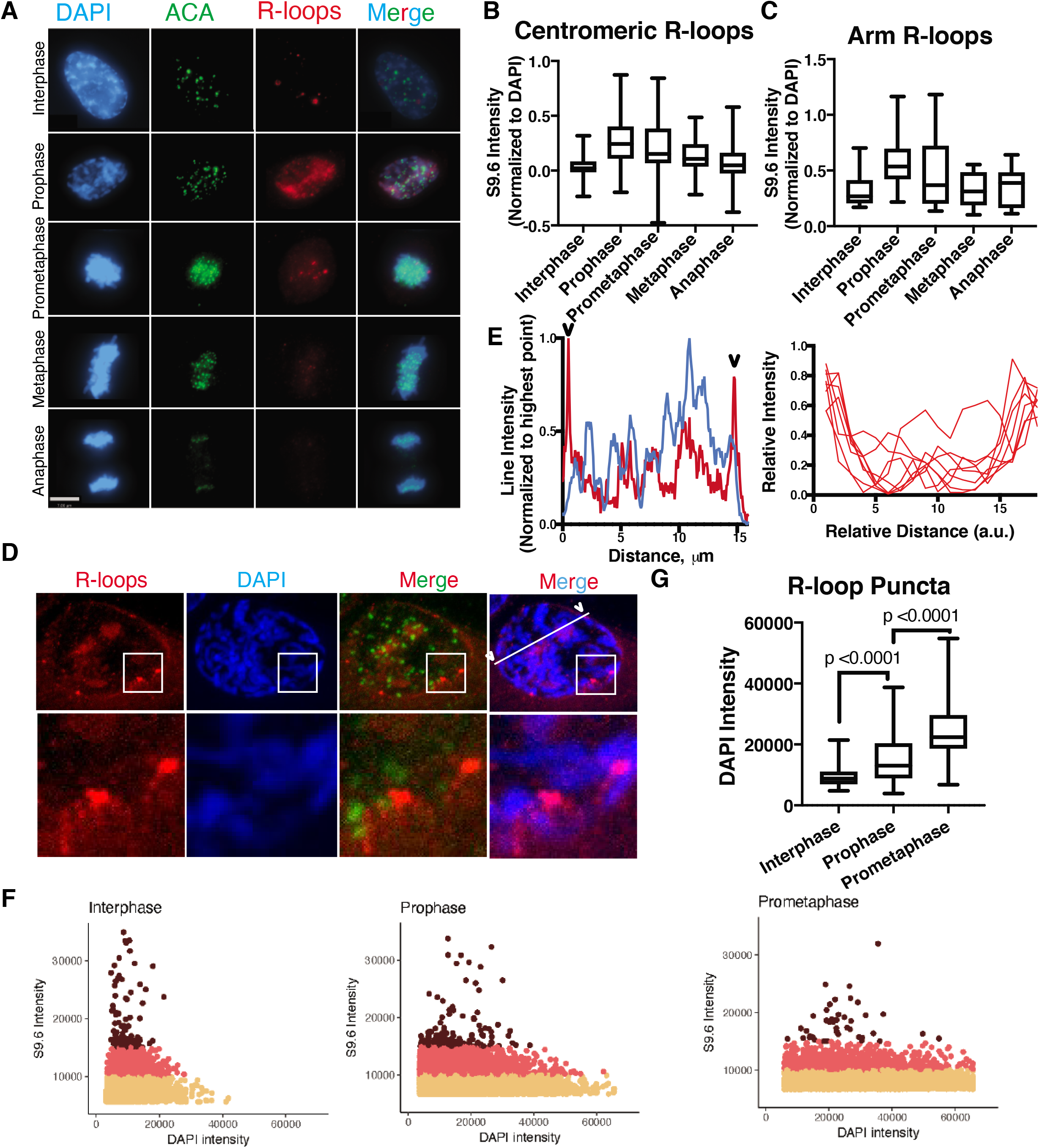
Mitotic R-loops are dynamic and associated with the nuclear periphery during prophase. A. Representative images from randomly cycling RPE1-T-REx cells after indirect immunofluorescence using antibodies against the human centromere (ACA), and R-loops (hybridoma S9.6). Images from one of 5 independent experiments. Scale bar, 7 μm. B-C. Quantification of S9.6 signal intensity overlapping with ACA (B) and DAPI with the ACA area subtracted (C), divided into interphase and 4 mitotic subsets, prophase, prometaphase, metaphase, and anaphase with 29, 50, 78, 38, and 46 cells measured respectively. Significance values are located in Figure 1—Figure supplement 1C, achieved by Kruskal-Wallis non-parametric ANOVA with post-test. D. Confocal microscopy images of a prophase nucleus. Arrowheads show where the line scan crosses the nuclear periphery. E. Left panel, relative intensity profile of line scan outlined in D. Right panel, 10 individual line scans from different prophase nuclei, scaled and overlaid. F. Scatter plots of intensity correlation between S9.6 and DAPI for a total of 3 nuclei at each phase, thresholded at 3000 a.u. and 5000 a.u. for S9.6 and DAPI respectively where S9.6 points were selected by the brightest points in a 0.1 μm 3-D rolling circle. S9.6 intensity was then subset into low, medium and high (tan, pink, and maroon). Maroon dots are quantified for the distribution of DAPI intensities in G. Significance values achieved by one-way parametric ANOVA with post-test. Interphase: n=117; Prophase: n=137; Prometaphase: n=42 puncta.

We analyzed the intra-nuclear location of prophase R-loops more precisely using confocal microscopy. Prophase R-loops were not observed uniformly across all chromosomes, and some chromosomes were depleted of R-loop foci. The most striking feature was that R-loops form along nuclear rims, which are classically associated with heterochromatin in interphase cells (Figure 2D, line scan in Figure 2E, right; additional prophase nuclei available in Supplemental figure 1). We noticed that R-loop foci localized to areas with low DAPI staining within the nuclei of prophase cells, suggesting they form before chromosomes are fully condensed. We quantified the brightest point of R-loop foci within the nuclei of cells. High R-loop foci were constrained to regions of relatively low DAPI intensity in each cell cycle state (Figure 2F, dark red points). Cells were continuously condensing chromatin over the cell cycle points measured. We noted that there was a gradual increase in DAPI intensity of the R-loop staining foci points as cells progress from interphase to prometaphase (Figure 2G). However, in each of the states, the high R-loop staining was always in low to moderate range of DAPI staining intensities (Figure 2F). This suggests that R-loops form on condensing chromatin and potentially represent an intermediate of chromatin condensation. Our data suggests that R-loops are most highly associated with condensing chromatin in early mitosis and associated with heterochromatin at the nuclear periphery.

### Aurora B kinase activity removes mitotic R-loops in randomly cycling cells

We performed immunofluorescence using S9.6 and anti-ACA antibodies in addition to DAPI on asynchronous RPE1-TREX cells, as in figure 2, treated with Aurora B kinase inhibitors ZM447439 (ZM) or AZD1152 (AZD) for 1 hour. AZD generated minimal increases in R-loop levels in interphase DNA and centromeres when normalized to ACA (p<0.01 and p<0.001 respectively), although this difference was not seen after ZM treatment (p=0.5 and p>0.9; Figure 3A-C). In contrast, both Aurora B inhibitors increased the levels of R-loops on mitotic chromosomes and centromeres (p< 0.01 and p<0.001 AZD-DAPI and AZD-ACA; p<0.001 and p<0.001 ZM-DAPI and ZM-ACA respectively; Figure 3A-C). There was no significant difference in ACA levels after treatment with either Aurora B inhibitors, validating its use as a normalization parameter (Figure 3D, ANOVA p-0.24). Thus, we were able to confirm that Aurora B has a role in removing R-loops from mitotic cells using both DRIP-Seq and immunofluorescence.

**Figure 3.**
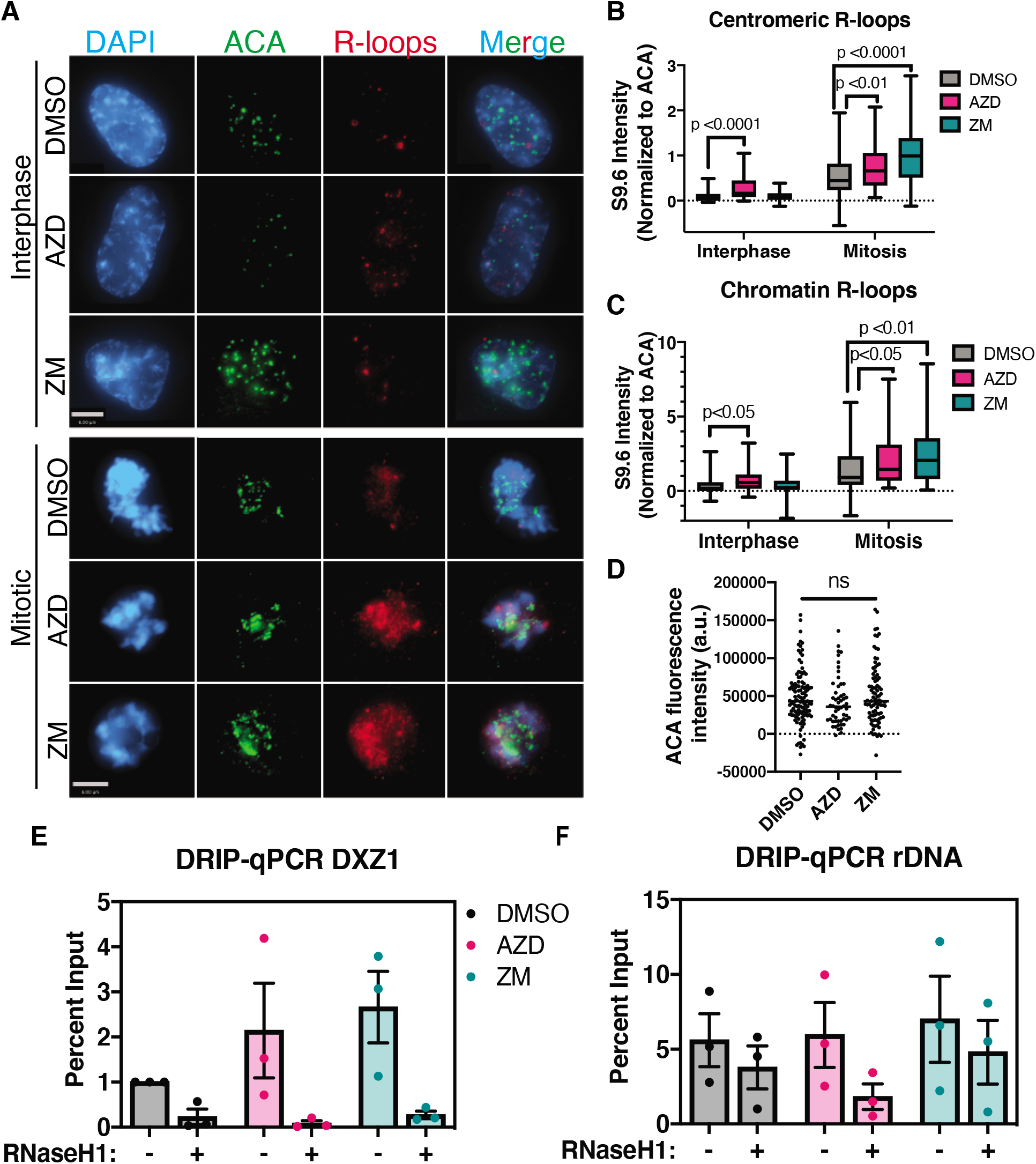
Aurora B activity promotes R-loop resolution in mitosis. A. Representative images of RPE1-T-REx cells treated with vehicle (DMSO), 500 nM AZD-1152, or 4 μM ZM-447439 for 1 hour. Images are from one of 4 experiments, quantified in B and C as the signal of S9.6 indirect immunofluorescence overlapping with signal of ACA (Centromeric R-loops, B) or DAPI (Chromatin R-loops, C), normalized to ACA signal. Kruskal-Wallis non-parametric ANOVA significance test performed to estimate a p-value. Interphase DMSO: n=27 cells; Interphase AZD: n=25 cells; Interphase ZM: n=28 cells; Mitotic DMSO: n=97 cells; Mitotic AZD: n=26 cells; Mitotic ZM: n=51 cells. D. ACA values were compared across samples to justify normalization to ACA, no significant differences were detected by Kruskal-Wallis ANOVA. E-F. DRIP-qPCR results from 3 independent experiments from DLD1 cells arrested in mitosis and treated with vehicle (DMSO), 500 nM AZD-1152, or 2 μM ZM-447439 for the last hour of the arrest. DXZ1 locus, X-chromosome primary a-satellite higher-order repeat array; rDNA assayed at 4 kb into the rDNA repeat.

We confirmed that Aurora B regulates centromeric R-loops as suggested by our DRIP-seq by performing DNA-RNA immunoprecipitation (IP) followed by quantitative PCR (DRIP-qPCR) utilizing primers to the X chromosome Higher Order Repeat of alpha-satellite (DXZ1 HOR) and to ribosomal DNA repeats (rDNA) in mitotically arrested cells. We found that treatment with either AZD or ZM for the last hour in cells stalled for 24 hours in colcemid caused at least a two-fold increase in R-loop accumulation at the DXZ1 HOR (Figure 3E-F). There is specificity for alpha-satellite sequences because Aurora B inhibitors had little effect on the accumulated R-loops at the rDNA locus. We conclude that Aurora B regulates R-loops repetitive regions during mitosis, particularly at centromeric alpha-satellites.

### R-loops promote Aurora B localization and activation

To determine the functions of R-loops during mitosis we created cells that express mCherry tagged *E. coli* RNaseH1 under a Tet-inducible promoter in HeLa-T-REx cells to generate an inducible system to deplete R-loops. We used clonal lines that expressed detectable levels of RNaseH1 within 8 hours of doxycycline-induced expression (Supplemental figure 2). Previous studies demonstrated that centromeric R-loops are required to activate the centromere pool of Aurora B but did not report a change to the amount of CPC (Kabeche et al., 2018). We localized the Aurora B kinase by immunofluorescence in order to determine if localization of Aurora B kinase is dependent upon R-loops. Induced RNaseH1 significantly depleted Aurora B intensity in mitotic cells compared to the same population of cells without RNaseH1 induction (Figure 4A and B) and compared with cells with induced expression of catalytically dead RNaseH1-2R mutant (Britton et al., 2014, Figure 4A and B). We stained chromosome spreads for S9.6 to observe R-loops and confirm that expression of RNaseH1-2R mutant does not deplete R-loops. Cells expressing the RNaseH1-2R mutant have R-loop staining throughout the chromosome (Supplemental figure 2).

**Figure 4.**
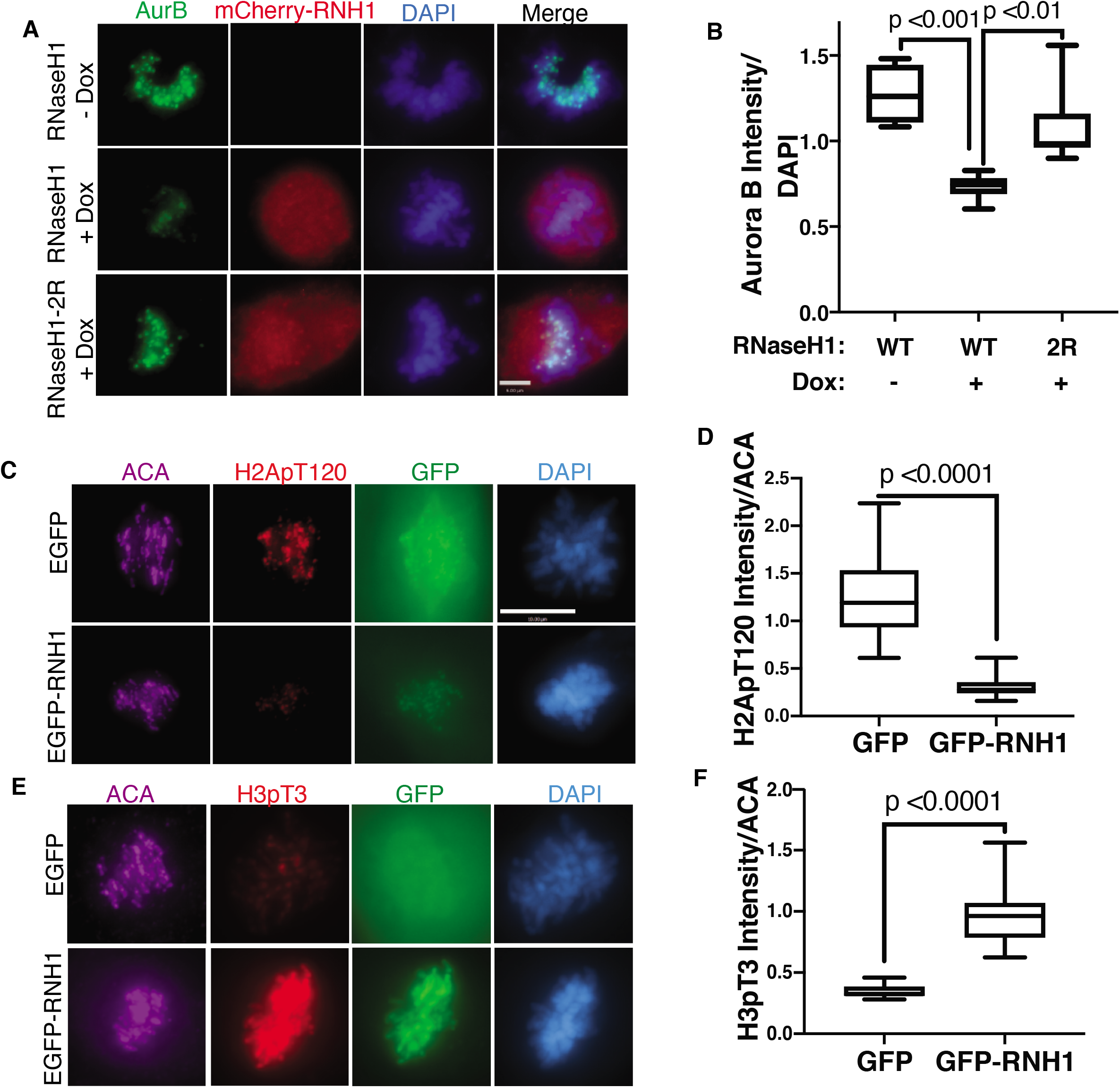
R-loop presence is necessary to localize Aurora B. A. Expression of a tet-inducible mCherry-RNaseH1 from *E.coli* and a catalytically dead mutant with point mutations D10R, E48R (2R) was induced within randomly cycling HeLa-T-REx cells and indirect immunofluorescence for Aurora B was assayed. Aurora B intensity normalized to DAPI intensity was quantified in B, with 10 cells measured in each; mean values and range are shown. Significance was determined by one-way ANOVA compared to WT cells. C-F. RPE1-T-REx cells infected with a constitutive EGFP or EGFP-hRNaseH1 construct, within a double thymidine block and release to allow for peak expression during the first mitosis. Cells were then subjected to indirect immunofluorescence for H2ApT120 and ACA (E-F), or H3pT3 and ACA (G-H). Signal of the histone mark normalized to ACA quantified in F and H, shown as median and range values. Significance values dictated by Mann-Whitney non-parametric tests.

We used a lentivirus to overexpress RNaseH1-EGFP in RPE1-TREx cells and then assessed intensity of Aurora B T-loop phosphorylation (pT232), as well as the signals that localize the CPC to the inner centromere, H3 pT3 and H2A pT120, in order to understand how R-loops control Aurora B activation and localization. Aurora B pT232 staining indicated a loss of auto-phosphorylated Aurora B in these cells after RNaseH1 overexpression (Supplemental figure 2) confirming that R-loops are involved in activation of the centromere pool of Aurora B (Kabeche et al., 2018) in addition to the pool on chromosome arms. We also observed a decrease in H2A pT120, one of the histone marks that serves as a localization signal (Figure 4C and 4D). We observed an increase in H3 pT3 throughout the chromosome, which explains the spread of Aurora B signal (Supplemental figure 2). This indicates that R-loops are required to reduce the H3 pT3 signal, and potentially Haspin kinase on chromosome arms. Together our data suggest that R-loops affect both histone marks that localize the CPC to drive the movements from arm chromatin to the inner centromere.

### The CPC interacts with R-loop regulators including RBMX on mitotic chromosomes

Aurora B has many functions in mitosis so it could have direct or indirect roles in resolving R-loops. We hypothesized that if the CPC had a direct role in R-loop resolution it should be bound to chromatin with other proteins that resolve R-loops. We developed an approach to rapidly purify the CPC bound to chromatin liberated from mitotic chromosomes based on the LAP dual affinity tag (Cheeseman and Desai, 2005). We established four HeLa cells lines expressing LAP-Aurora-B, LAP-Borealin, LAP-Survivin or LAP only (Supplemental figure 3). The LAP-Survivin, and LAP-Borealin proteins localized to centromeres in mitosis (Supplemental figure 3). We rapidly isolated mitotic chromatin by a clarification centrifugation followed by pelleting mitotic chromatin. Chromatin was liberated by micrococcal nuclease treatment producing ladders that ranged from mononucleosomes to hexameric nucleosomes. The CPC from mitotic chromatin was tandem-affinity purified and the bound proteins were analyzed by MudPIT (Supplemental Table 1, Supplemental figure 3), which identified a total of 111 proteins (Figure 5A). All three CPC complex members purified at least one other CPC member and no CPC proteins were identified in the LAP control preps. We identified Topo IIα, Kif20a/MKLP2 and HP1β which have been previously shown to interact with the CPC (Coelho et al., 2008; Gruneberg et al., 2004; Kang et al., 2011; Morrison et al., 2002). The top three GO keywords for proteins identified were Phosphoproteins, Ribonucleoproteins, and RNA-binding, consistent with the fact that RNA has a major role in CPC activity (Figure 5B, Blower, 2016; Jambhekar et al., 2014). The majority (75%) of the RNA binding proteins that interact with the CPC were also purified in a S9.6 IP (the R-loop interactome, Figure 5B, Cristini et al., 2018). 35% of these proteins (11 proteins) were also identified as Aurora kinase substrates in phosphoproteomic screens. The fact that the CPC interacts with and phosphorylates R-loop proteins strongly suggests that the CPC has a direct role in controlling R-loops and all of these proteins are potential regulators of mitotic R-loops.

**Figure 5.**
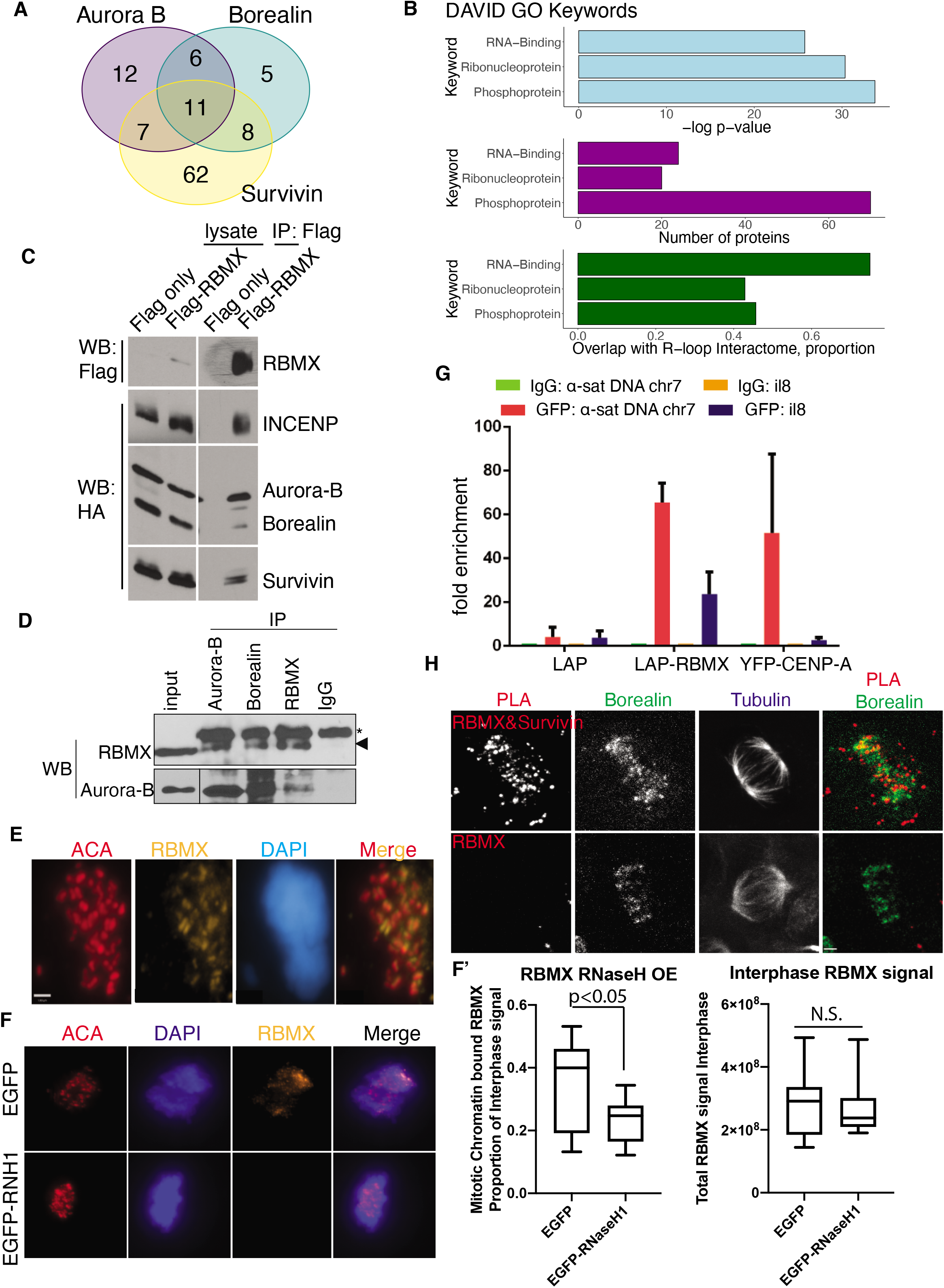
RBMX associates with the CPC and R-loops. A. MudPiT analysis of LAP-tagged Aurora B, Borealin, and Survivin precipitation many of the same peptides, not observed in the background LAP only precipitation. B. DAVID GO keywords of all proteins precipitated in A, top three are phosphoprotein, ribonucleoprotein, and RNA-binding protein. Light blue, negative log p-value as identified by DAVID; purple, number of proteins associated with each keyword; green, proportion of the total proteins associated with each keyword that also appears in the R-loop interactome published by (citation). C. Co-immunoprecipitation of tagged proteins from HEK293T cells. Flag-tagged RBMX or Flag alone was co-expressed with HA-tagged CPC members, and all four CPC members can be observed to be co-immunoprecipitated with RBMX specifically. D. Co-immunoprecipitation of endogenous protein from HeLa-T-REx cells. Immunoprecipitation of endogenous Aurora B, Borealin, and RBMX but not non-specific IgG can be observed to precipitate RBMX and Aurora B. E. Indirect immunofluorescence after extraction of unbound proteins prior to fixation in RPE1-T-REx cells shows RBMX localization to the centromere, as marked by ACA. Scale bar, 1.6 μm. F. Degradation of R-loops by overexpression of EGFP-hRNaseH1 in RPE1-T-REx cells leads to loss of RBMX bound to chromatin. Representative images of prometaphase cells in the left panel. F’ Quantification of chromatin bound RBMX in mitotic cells, expressed as a proportion of measured nearby interphase cells as the maximal proportion of RBMX protein that could remain bound to chromatin, which is shown in the right panel. Graphs show mean and range values for the 15 mitotic and 15 paired interphase cells, P-value calculated by unpaired two-tailed t-test. G. ChIP-qPCR of LAP, LAP-RBMX, or YFP-CENP-A to the a-satellite array of chromosome 7 or control locus Il8. Values are expressed as fold enrichment of IgG control IP from the same cells at the same locus. Graph shows the results of 2 ChIP experiments. H. Proximity ligation assay (PLA) of endogenous RBMX and Survivin from HeLa-T-REx cells. Borealin indirect immunofluorescence was also performed as a localization control for the CPC.

We initially focused on RBMX because it was identified in the R-loop interactome and is required for centromeric cohesion (Matsunaga et al., 2012). We confirmed that RBMX interacts with the CPC by co-IP (Figure 5C, D). The bulk of RBMX is cytoplasmic in mitosis (Matsunaga et al., 2012), but after extracting soluble proteins in RPE1-T-REx cells the majority of the remaining RBMX colocalized with ACA (Figure 5E) and this localization is dependent upon R-loops (Figure 5F and F’). We generated a cell line expressing LAP-RBMX to confirm whether RBMX is enriched on centromeres. LAP-RBMX is greatly enriched at centromeres as measured by Chromatin Immunopreciptitation (ChIP) using primers against α-satellite DNA (Figure 5G). We confirmed the localization by Proximity Ligation Assay (PLA), which measures the proximity of the GFP of the LAP tag to the CPC subunit Survivin in LAP-RBMX expressing cells. Cells were co-stained with antibodies to Borealin and tubulin to generate fiducial marks on inner centromeres and the spindle. PLA signals were found adjacent to Borealin at centromeres of metaphase cells (Figure 5H) and there was little signal if the Survivin antibody was not added to the reaction as a negative control. We conclude that RBMX is recruited to centromeres by R-loops where it interacts with the CPC.

We tested whether RBMX has a function in mitotic R-loop biology. We depleted RBMX by shRNA and co-stained the cells with anti-Aurora-B, ACA, and S9.6 antibodies to determine whether RBMX regulates Aurora-B and R-loops. RBMX protein levels were reduced as evaluated by western blot but the levels of Aurora-B, Survivin, and a set of cohesion regulators were unaffected by depletion of RBMX, suggesting RBMX does not interfere with transcription or protein stability of the CPC (Supplemental figure 4). Depleting RBMX affects Aurora B localization to centromeres as the amount of Aurora-B at inner centromeres of prometaphase cells was significantly reduced (Figure 6A-D). R-loop levels were significantly increased across chromatin in prometaphase cells, consistent with a loss of Aurora B activity (Figure 6A-B, Figure 3). We also saw reduction in the levels of the Borealin subunit of the CPC and T-loop phosphorylation of Aurora B kinase (Supplemental figure 4). The reduction of CPC was seen with a second shRNA that reduced RBMX protein levels (Supplemental figure 4), in two cell types (RPE-T-REx cells and HeLa-T-REx cells, Figure 6A-D) and expression of shRNA resistant LAP-RBMX restored centromeric Aurora-B levels to cells treated with shRNA against RBMX (Figure 6C, D). Thus, RBMX is required for centromeric accumulation of Aurora B and mislocalizing Aurora B is not an off-target effect of shRNA expression. RBMX is required for the histone marks that target the CPC to the centromere, but these marks can be restored by forced targeting Aurora B kinase activity to centromeres (Supplemental figure 5), demonstrating that RBMX is not required to generate the histone marks, but they are reduced in cells that are depleted of RBMX because Aurora B is missing. In conclusion, these data suggest that the RNA binding protein, RBMX, is recruited to centromeres by R-loops, where it recruits Aurora B to resolve R-loops.

**Figure 6.**
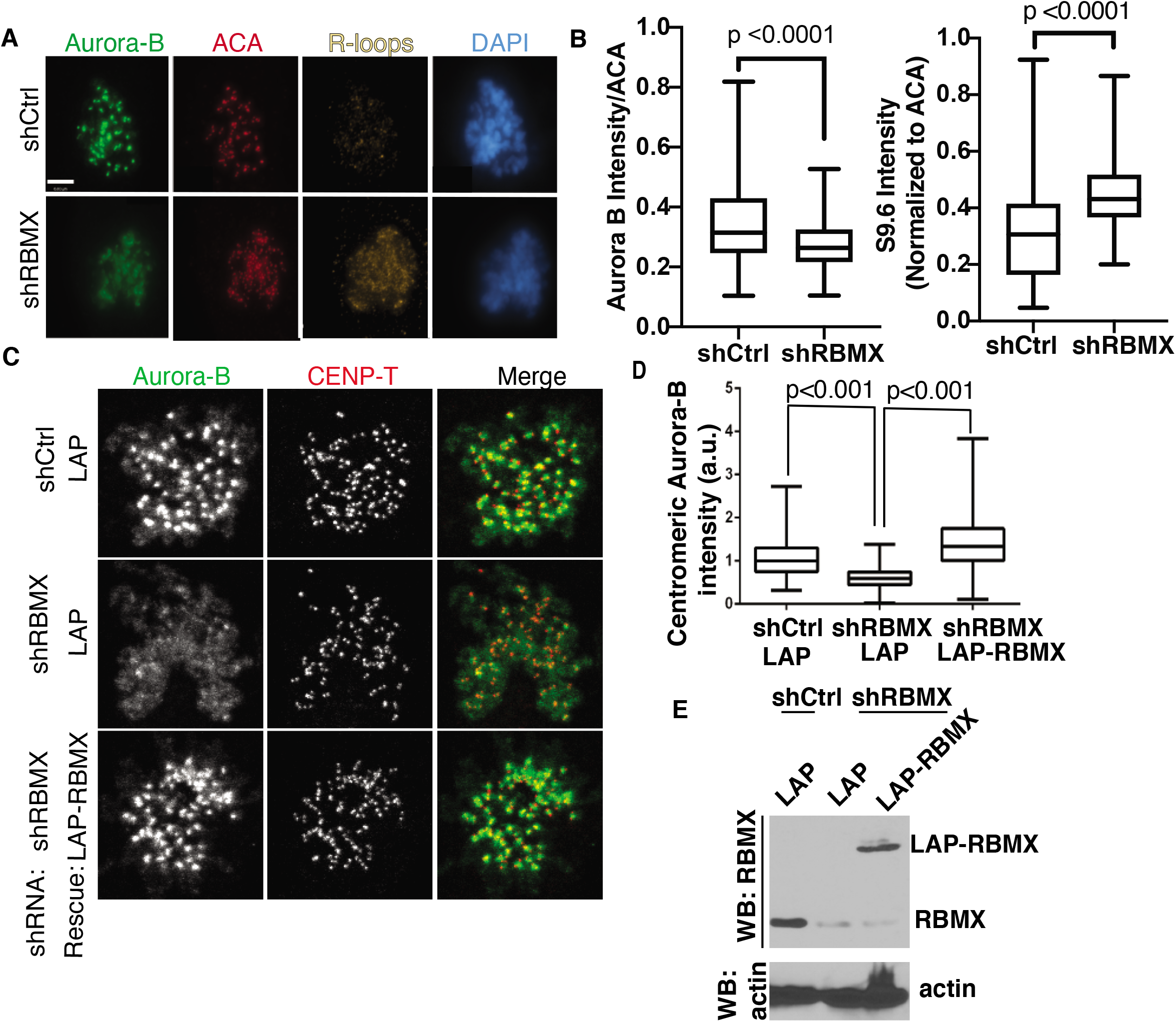
RBMX is necessary to resolve R-loops and localize Aurora B. A-B. shRNA knockdown of RBMX in RPE1-T-REx cells results in loss of Aurora B localization, as well as an increase in prevalence of S9.6 immunofluorescence signal. A. Representative images from one of 3 experiments, quantified in B, of the loss of Aurora B immunofluorescence intensity and gain of S9.6 fluorescence intensity. Graphs in B show median and range values for normalized fluorescence intensity, n=22 cells and 15 cells respectively. P-values estimated by Mann-Whitney non-parametric tests. C-E. loss of Aurora B after shRNA knockdown of RBMX in HeLa-T-REx cells can be rescued by expression of LAP-RBMX. C. Representative images of immunofluorescence of Aurora B and CENP-T from 2 experiments. D. Quantification of centromeric Aurora B, normalized to CENP-T immunofluorescence signal. P-value determined by one-way ANOVA, n=17, 12, 18 respectively. E. Western blot showing RBMX expression after shRNA knockdown and re-expression of LAP-RBMX.

### R-loops and RBMX are required to maintain centromeric cohesion and the role of RBMX in centromeric cohesion is to recruit Aurora B and Sgo1

We have shown that RBMX is localized by R-loops (Figure 5) and it has been previously published that RBMX is required for centromeric cohesion (Matsunaga et al., 2012) To identify the function of mitotic R-loops, we tested whether loss of R-loops affects cohesion. We expressed RNaseH1 and RNaseH1-2R in HeLa-T-REx cells and performed mitotic spreads and quantified premature chromatid separation (PCS, Figure 7A). Centromeric cohesion was lost in cells overexpressing RNaseH1 but not in cells expressing RNaseH1-2R (Figure 7B). To test whether RBMX targets the CPC to generate centromeric cohesion, we depleted RBMX and induced CENP-B-INCENP (CB-INCENP) fusion protein expression to forcibly target the CPC to the centromere via CENP-B binding. RBMX depleted cells showed a PCS phenotype, however PCS was dramatically decreased when these cells were induced to express the CB-INCENP (Figure 7C). The loss of centromeric cohesion was restored by targeting the Aurora B to centromeric chromatin. We conclude that R-loops are required to maintain centromeric cohesion by RBMX-dependent recruitment of the CPC.

**Figure 7.**
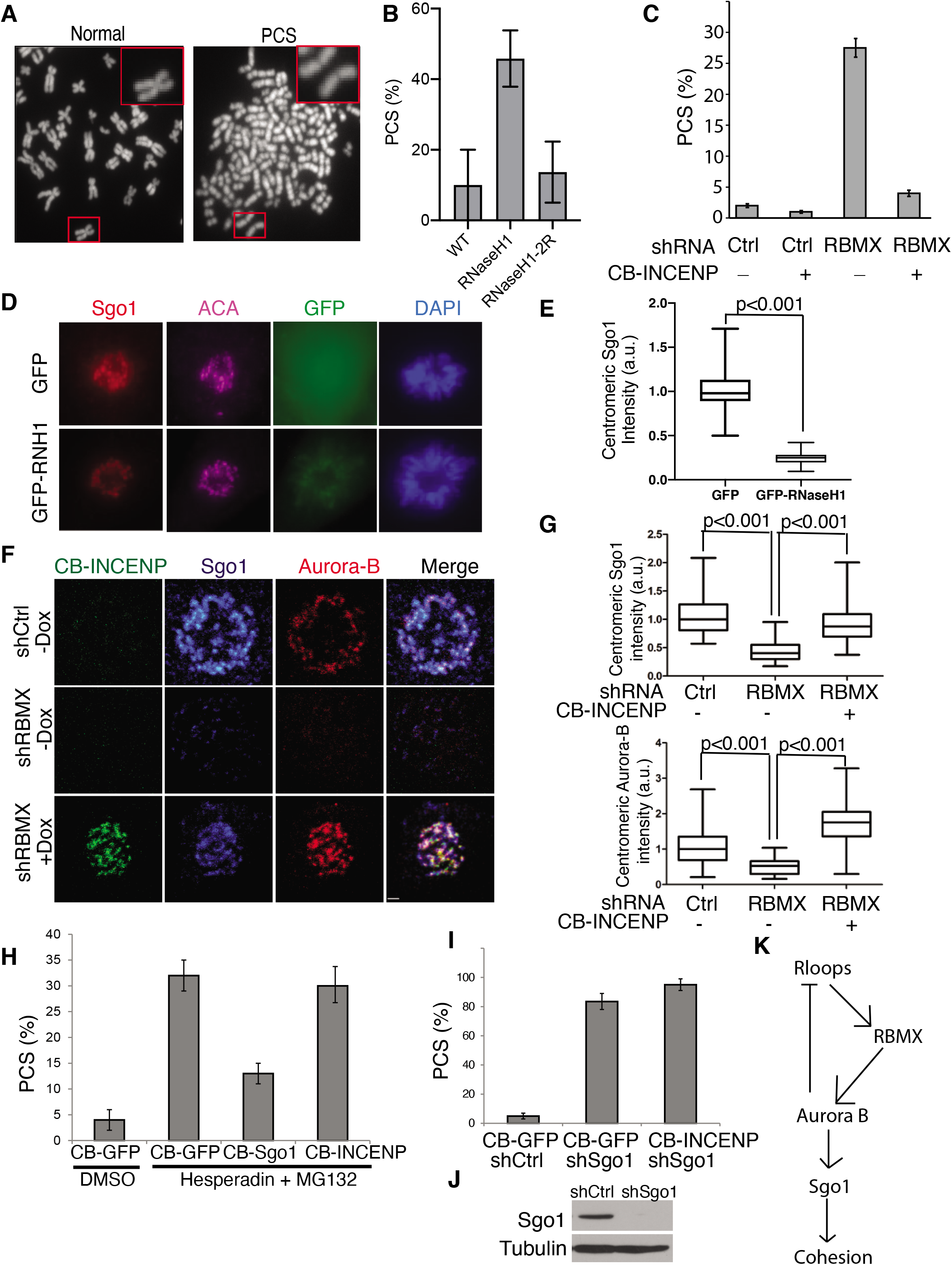
R-loops and RBMX recruit Aurora B to recruit Sgo1 and maintain centromeric cohesion. A. Representative images of normal spreads and cells with premature chromatid separation (PCS). B. Quantification of percent PCS in HeLa-T-REx cells overexpressing *E. coli* RNaseH1 or catalytically dead RNaseH1-2R mutant from 4 replicate experiments, with 10 fields imaged from each cell type in each experiment. C. Quantification of percent PCS in HeLa-T-REx cells expressing shRNAs that are either non-targeting (shCtrl) or against RBMX and a tet-inducible CENP-B DNA binding domain fused to INCENP. 3 replicate experiments. D. RPE1-T-REx cells overexpressing GFP-hRNaseH1 or GFP empty vector, stained using indirect immunofluorescence for Sgo1 and ACA. Quantified in E, p-value determined by unpaired two-tailed t-test. F. Cells in C stained for immunofluorescence of Sgo1 and Aurora B. Quantified in G for Sgo1 (top) and Aurora B (bottom) intensity, normalized to the level of fluorescence in control shRNA cells without CENP-B-INCENP construct expression. P-values determined by one-way ANOVA. H. PCS assay for HeLa-T-REx cells expressing CENP-B DNA binding domain fused with GFP, Sgo1, or INCENP and either treated with vehicle (DMSO), or Aurora B inhibitor Hesperadin at 100 nM with MG132 to prevent escape from mitotic arrest. 3 replicate experiments. I. PCS assay for HeLa-T-REx cells expressing non-targeting shCtrl or shSgo1, and CENP-B DNA binding domain fused to either GFP or INCENP. 3 replicate experiments. J. Western blot for cells expressing non-targeting shCtrl or shSgo1 and blotted for Sgo1. K. Schematic representation of the results of the epistatic experiments presented in this work.

Sgo1 is a key regulator of centromeric cohesion, and Aurora B activity is required to localize Sgo1 to inner centromeres (Dai et al., 2006; van der Waal et al., 2012). We explored the relationship of the R-loops with the localization of Sgo1 to determine the mechanism by which RBMX and R-loops maintain centromeric cohesion. Expressing RNaseH1 reduced the centromeric levels of Sgo1 significantly (p<0.0001, Figure 7D, E). Depletion of RBMX similarly reduced centromeric levels of Sgo1, and this could be recovered by forced targeting of the CPC by CB-INCENP (Figure 7F, G). We conclude that RBMX targets the CPC to recruit Sgo1.

### The pool of the CPC recruited by RBMX protects cohesion by recruiting Sgo1

Our finding that targeting the CPC to centromeres can reverse the PCS that was induced by RBMX depletion suggested that this pool of the CPC is involved in protecting centromeric cohesion. It is difficult to assay a role for the CPC in centromeric cohesion because the CPC is required to remove cohesin from non-centromeric regions during prophase (Giménez-Abián et al., 2004; Losada et al., 2002; Nishiyama et al., 2013). Thus, even if cells depleted of Aurora-B lost centromeric cohesion the chromatids would remain cohesive through the chromosome arms. Arm cohesion is removed but centromeric cohesion is still preserved when human cells are arrested with microtubule poisons for 2-3 hours (Giménez-Abián et al., 2004). We therefore modified an assay that previously showed that the CPC is required to maintain centromeric cohesion (Tanno et al., 2010). We generated a population of mitotic cells by initially synchronizing cells in S-phase by double thymidine block and then ten hours after release from thymidine we treated cells with colcemid for 3 hours to allow the cells to arrest in mitosis and lose arm cohesion. We then treated the cells with Aurora-B inhibitors for an additional two hours and measured PCS by mitotic spreads (Figure 7A). Forty percent of mitotic cells showed PCS when treated with 0.1μM Hesperadin while only 2% PCS occurred when treated with DMSO (Supplemental figure 6). There was a dose dependent increase of PCS when cells were treated with Hesperadin. Treatment of cells with ZM, a structurally different Aurora-B kinase inhibitor, also promoted PCS, demonstrating that Aurora-B activity is required for centromeric cohesion (Supplemental figure 6). To rule out the possibility that separase is prematurely activated under our experimental conditions; we treated cells with MG132, a proteasome inhibitor. We observed similar loss of cohesion in the presence or absence of MG132 (Supplemental figure 6).

To circumvent the concern that the CPC’s role in cohesion is a consequence of prolonged mitotic arrest we complemented our findings by depleting Aurora-B by shRNA in a HeLa line that stably expressed LAP-CENP-A, visualized mitotic chromosomes by chromosome spreads, and measured the inter-kinetochore distance (Supplemental figure 6). All chromosomes maintained cohesion consistent with the requirement of the CPC to remove arm cohesion. However, there was a 36% increase of the inter-kinetochore distance on Aurora-B depleted chromosomes, supporting the hypothesis that Aurora-B is required for centromeric cohesion.

We determined the epistatic relationship between Aurora-B and Sgo1 in centromeric cohesion regulation. Forced targeting of Sgo1 to centromeres using a Cenp-B Sgo1 fusion protein partially rescued Hesperadin-induced PCS (Figure 7H), suggesting Sgo1 is downstream of CPC mediated protection of centromeric cohesion. In contrast, forced targeting of INCENP to centromeres did not rescue Sgo1 depletion induced PCS (Figure 7I), consistent with Aurora-B being upstream of Sgo1. Finally, targeting INCENP to inner centromeres also did not rescue hesperidin-induced PCS (Figure 7H), indicating the kinase activity of Aurora-B is required for protection of centromeric cohesion. These data suggest that R-loops recruit RBMX, which in turn recruits the CPC to recruit Sgo1 and maintain centromeric cohesion (Figure 7J). In addition, Figure 3 and 4 suggest Aurora B also reduces centromeric R-loops, suggesting a self-limiting feedback loop.

## Discussion

R-loops are emerging as critical regulators of interphase chromatin, but much less is known about their function in mitosis. Surprisingly, we found that the R-loops were higher in regions of intermediate chromatin density in prophase cells than interphase cells, suggesting that R-loops are formed during chromosome condensation. R-loops decline during prometaphase and metaphase until they reach interphase levels at anaphase. We explored the mechanism that cells use to resolve mitotic R-loops and found that this process was dependent upon Aurora B kinase activity. Purification of the CPC from mitotic chromosomes identified 32 proteins that were also found in purifications of R-loops. We verified that one of these, RBMX, is required to remove mitotic R-loops. Finally, we explored the function of mitotic R-loops and found they localize the CPC to centromeres to maintain sister chromatid cohesion.

We have established that R-loops form during mitosis in distinct locations from those in interphase cells and there are active mechanisms to resolve them. In addition, we show that mitotic R-loops are required for mitotic chromosome cohesion and this is mediated by R-loops recruiting the CPC to inner centromeres. The CPC is both recruited by R-loops and resolves these R-loops demonstrating feedback control. In addition to the CPC, we have identified a set of potential regulators of mitotic R-loops but purifying the CPC from mitotic chromatin. We have confirmed one of these proteins RBMX is both required to recruit the CPC to inner centromeres and resolve R-loops. Importantly, loss of RBMX is also required for centromeric cohesion.

Our data are consistent with a recent paper that suggested that R-loops exist in mitotic centromeres where they recruit the ATR kinase to activate centromeric Aurora B (Kabeche et al., 2018). Our findings support this model and extend it by showing that R-loops are required to localize Aurora B to inner centromeres. Another study suggested that the phosphorylation of Histone H3 on Serine 10 is found at R-loops containing chromatin in yeast (Castellano-Pozo et al., 2013a). Aurora B is the writer of this histone mark, and our finding that Aurora kinase activity is required to resolve R-loops in human cells suggests that R-loop regulation is a conserved function of the histone H3 S10 phosphorylation. A previous study also showed R-loop association with H3 lysine 9 dimethylation, a marker of condensed chromatin. This is consistent with our demonstration that R-loops are highest in prophase, when chromosomes condense.

We employed DRIP-seq assays to identify the genomic positions of R-loops in populations of cells arrested in mitosis and the genomic loci of R-loops regulated by Aurora B. Mitotic R-loops were depleted from gene bodies, but highly enriched at repetitive DNA, most notably alpha-satellite and SAR sequences. Enrichment at alpha-satellite is consistent with roles of Aurora B in centromeric regulation and cohesion. SARs were associated with loop regions of chromatin in older studies (Mirkovitch et al., 1987; Strissel et al., 1996) but their function is still poorly understood; it is also notable that the genomic loci of SAR repeats are within pericentric DNA (Hubley et al., 2016). We identified three SAR binding proteins SAF-A, SAF-AL and SAF-B in purifications of CPC bound to mitotic chromatin. The connection between SARs and the CPC from two independent unbiased methods suggests this to be an important connection. We speculate that R-loops and the CPC act at the base of the loops of condensing chromatin. This is supported by the association of the CPC with condensin and Topo II, which are localized to the base of chromatin loops in mitotic chromosomes. It is also consistent with a recent study that suggested that R-loops template the base of chromatin loops of maize centromeres (Liu et al., 2020).

The source of mitotic R-loops is a second area of study suggested by our results. Interestingly, condensin acts at the base of chromosome loops and can generate positive supercoiling (Bazett-Jones et al., 2002; Kimura and Hirano, 1997; Kimura et al., 1999). Topoisomerases are known to work with condensin to relieve topological strain (Baxter et al., 2011), and the activity of topoisomerases could expose single-stranded DNA and allow RNAs to hybridize, especially within highly repetitive sequences. R-loops have been reported at sites of negative supercoiling (Stolz et al., 2019), and sites of topoisomerase activity (Drolet et al., 1995; El Hage et al., 2010). R-loops might also be a result of condensing heterochromatic regions as the cell moves from interphase to mitosis. H3 phosphorylation on S10 has been shown to displace Heterochromatin Protein 1 (HP1, Hirota et al., 2005) which binds H3 lysine 9 methylation. We found that R-loops are highest at the nuclear periphery in prophase cells, and that the largest proportion of DRIP peaks existed at repetitive sequences which are marked by H3 lysine 9 methylation in interphase.

We purified the CPC from mitotic chromatin to gain insight on how it would resolve R-loops and found a pool of proteins in this proteome that associated with purified R-loops (Cristini et al., 2018). We initially focused on RBMX because it had been shown to regulate cohesion in a Sgo1-dependent manner (Matsunaga et al., 2012) and our epistasis experiments suggest a pathway whereby R-loops recruit RBMX to recruit Aurora B to resolve R-loops (Figure 7I). We believe that our preparation of proteins associated with the CPC on mitotic chromatin and particularly those that also interact with R-loops will be a rich source of future studies and will help us understand the precise nature of the generation and resolution of R-loops in mitosis.

Our work sheds light on a number of questions associated with mitotic R-loops. First, we established that there is a specific population of R-loops that arise and are resolved within the course of mitosis that are distinctly distributed within the genome from interphase R-loops. These R-loops are preferentially enriched within repetitive sequences, including centric and pericentric repeats. We have also identified a number of proteins involved with the regulation of mitotic R-loops, including Aurora B kinase and a number of other chromatin and RNA regulators. Aurora B kinase has an active role in limiting the formation of repetitive R-loops, and we hypothesize that this is through phosphorylation of substrates found within the pool of R-loop regulators. Although R-loops likely have multiple roles in mitosis, we have identified a pathway linking R-loops to regulation of centromeric cohesion, demonstrating that this regulation is crucial to maintenance of mitotic fidelity and may provide a mechanism for the increase of lagging chromosomes found in cells overexpressing RNaseH1 (Kabeche et al., 2018). Overall, this work links R-loops to major mitotic regulator Aurora B and gives a mechanism for the observation that RNA can regulate Aurora B and centromeres through *cis*-activity in human cells(Grenfell et al., 2016; Perea-Resa and Blower, 2017).

## Materials and Methods

### Cell Culture

RPE1-T-REx cells were generated from RPE1-hTERT (ATCC) cells using the T-Rex system (Thermo Scientific) plasmid by transfection and selecting for stable integration using Blasticidin. These cells were cultured using DMEM/F12 1:1 (Gibco) supplemented with 10% (vol/vol) FBS (Gibco), penicillin and streptomycin. HeLa-FRT-T-REx cells were generated using the Flp-in T-REx system (Thermo Scientific) and were a gift from the Dan Foltz lab. HEK-293T cells (ATCC) and HeLa-FRT-T-REx cells were cultured in DMEM (Gibco) supplemented with 10% (vol/vol) FBS, penicillin and streptomycin. DLD1 cells were a gift from the Michael Guertin lab, and were cultured in RPMI (Gibco) media supplemented with 10% (vol/vol) FBS, penicillin and streptomycin. All cells were grown in a humidified chamber at 37 ^0^C in the presence of 5% CO_2_.

### Stable cell lines generation

HeLa-T-REx-RNaseH1 and RNaseH1^D10R, E48R^ (2R) were created by transfecting pICE-RNaseH1-WT-NLS-mCherry and pICE-RNaseH1-D10R-E48R-NLS-mCherry (a gift from Patrick Calsou, Addgene plasmids #60365 and #60367 Britton et al., 2014) using Lipofectomine 2000 (Invitrogen) and selecting using 1 μg/ml Puromycin for 2 weeks. Clonal lines were created by plating at very low density and selecting 30 colonies of each and selecting for colonies that were mCherry negative in absence of doxycycline and induced detectable fluorescence within 4 hours of addition of 1 μg/ml doxycycline.

The LAP tag from pIC113 vector (Cheeseman and Desai, 2005) was subcloned into pcDNA5.0/FRT vector (Invitrogen) to make the pcDNA5.0/FRT-LAP-N vector. Full length cDNAs of Aurora-B, Borealin, and Survivin were cloned into pcDNA5.0/FRT-LAP-N vector to make LAP-Aurora-B, LAP-Borealin and LAP-Survivin constructs respectively. These LAP tagged contructs were co-tansfected with pOG44 (Invitrogen) into Flp-In HeLa T-REx cells. Stable lines expressing these constructs were created by selection with hygromycin (200μg/ml, Invitrogen) for two weeks.

Full-length cDNAs of Aurora-B, Borealin, INCENP and Survivin were cloned into pcDNA3.0-HA vector to make HA-tagged Aurora-B, Borealin, INCENP and Survivin respectively. DLAP and DHA destination vectors were made based on pcDNA5.0/FRT-LAP-N and pcDNA3.0-HA vectors respectively. The cDNA of RBMX were obtained from the human ORFeome collection (V5.1) and was cloned into DLAP and DHA destination vectors using gateway cloning technology (Invitrogen) to make LAP-RBMX and HA-RBMX. The LAP tag from pIC113 vector (Cheeseman and Desai, 2005) was subcloned into pcDNA5.0/FRT/TO vector (Invitrogen) to make the pcDNA5.0/FRT/TO-LAP-N vector. CB-INCENP-GFP vector is a kind gift from M.A Lampson (Liu et al., 2009). We amplified CENP-B 1-158 (CB) by PCR and clone it into pcDNA5.0/FRT/TO-LAP-N vector using Not I and BamH I sites, which removes sequence encoding S peptide and leaves GFP sequence intact. This generated pcDNA5.0/FRT/TO −CB-GFP vector (CB-GFP). The full length cDNA sequence of Sgo1 and the cDNA sequence encoding INCENP aa47-aa917 were cloned into pcDNA5.0/FRT/TO-CB-GFP vector to generate the pcDNA5.0/FRT/TO-CB-GFP-Sgo1 (CB-Sgo1) and pcDNA5.0/FRT/TO-CB-GFP-INCENP 47-917 (CB-INCENP). The stable lines constitutively expressing LAP-RBMX or inducibly expressing CB-GFP, CB-Sgo1 or CB-INCENP were made by co-transfecting these constructs with pOG44 (Invitrogen) into Flp-In HeLa T-REx cells and selection with hygromycin (200μg/ml, Invitrogen) for two weeks.

The human lentiviral shRNAmir pGIPZ constructs were obtained from Open Biosystems and grown and purified according to their protocol. The targeting sequences of the shRNAs used in this study are listed in Table 3. To package virus, 1.5 x10^7^ HEK-293T cells were co-transfected with 18 μg pGIPZ plasmid, 6 μg pMD2G plasmid, and 12 μg psPAX2 plasmid. Medium were replenished 24 hours after transfection and supernatants containing virus were collected and filtered through 0.2μm filters 48 hours after transfection. Cells were infected with virus in the presence of 8μg/ml polybrene (Sigma).

To create RPE1-T-REx EGFP, EGFP-RNaseH1, and EGFP-RNaseH1^D201N^ cells, EGFP-hRNaseH1 or EGFP alone was cloned into pDONR-221 via Gateway cloning (Invitrogen) and then recombined into pLX-304 (Gift from David Root, Addgene plasmid # 25890 Yang et al., 2011). Site-directed mutagenesis on pDONR221-EGFP-hRNaseH1 using primers in Table 3 and then recombined into pLX-304. Virus was packaged as above. Double thymidine synchronized cells were infected with viral supernatant without polybrene upon release from the first thymidine stall and again upon second thymidine stall to achieve 100% infection and expression in the cell cycle following second thymidine release.

### Immunoblotting and immunoprecipitation

For immunoprecipitation, HEK-293T Cells were co-transfected with plasmids encoding HA-tagged −Aurora-B, −Borealin, −INCENP, −Survivin and Flag-tagged, −RBMX. Forty-eight hours after transfection, cells were synchronized to mitosis with 100ng/ml colcemid for 16 hours. Cells were lysed in lysis buffer (250mM NaCl, 50mM Tris-HCl, 5mM EDTA, 0.5% NP-40, 1 mM DTT, 20mM Beta-glycerophosphate, 50mM NaF, 1mM Sodium orthovanadate, 1x protease inhibitors cocktail (Roche) and sonicated with cell disruptor for 30 cycles with 30 seconds on and 30 seconds off at 4 °C. The whole cell extracts were cleared by centrifugation at 16000g for 20 minutes and the supernatants were subjected to immunoprecipitation with EZView anti-flag beads (Sigma) for 4 hours at 4 °C. The beads were washed three times with lysis buffer. The bound proteins were resolved on 6-18% SDS-PAGE gel and blotted with antibodies as indicated.

### Immunofluoresence microscopy

HeLa T-REx cells were seeded onto coverslips coated with poly-L-Lysine (Sigma) one day before staining. The cells were co-fixed with 2% paraformaldehyde, PHEM buffer (60 mM Pipes, 25 mM Hepes, 10 mM EGTA, and 4 mM MgCl2, pH 6.9), and 0.5% Triton-X 100 for 20 minutes at room temperature. Cells stained with RBMX were pre-extracted with 0.5% Triton-X 100 for 2 minutes prior to fixation as above. After washing with PBS for three times, cells were blocked with 1% BSA for 30 minutes. Immunostaining was performed with primary antibodies (Table 1) at the indicated dilution for 1 hour at room temperature. After washing three times with PBS, cells were incubated with fluorescent secondary antibodies (Jackson ImmunoResearch). After washing two times with PBS, the cells were counterstained with 0.5μg/ml DAPI for 5 minutes. After two more washes with PBS, the coverslips were mounted onto slides using ProlongGold Antifade (Invitrogen) and sealed with nail polish. Image acquisition was performed as described previously (Banerjee et al., 2014), or on a Zeiss 880 confocal microscope in the UVA Advanced Microscopy Facility (Figure 1). Images were processed and analyzed using Volocity (V6.3, PerkinElmer). To quantify fluorescence levels at centromeres, we used a volume thresholding algorithm to mark all centromeres on the basis of ACA or CENP-A staining in projected images. To eliminate the size difference of each marked centromere, the sum of the fluorescence intensity was divided by the voxel volume to obtain the value of fluorescence intensity per volume. To quantify fluorescence over chromatin, a volume thresholding algorithm was applied to the DAPI signal and volume normalized. After background subtraction, we calculated the intensity/volume values for each channel. These values were normalized against the corresponding ACA or CENP-A intensity/volume. When cells were not stained with a centromere marker (ACA or CENP-A), we marked centromeres based on Bub1, Aurora-B or Sgo1 staining using a volume thresholding algorithm. These values were plotted using Prism (GraphPad) and the statistical significance was determined by the appropriate statistical test for the data, defined by normality and number of comparisons. For Box-and whisker plots, central lines indicate medians and whiskers are from minimum to maximum (range 0-100 percentile). For PCS assay, chromosome spreads were performed where cells were treated with 100ng ml-1 colcemid were trypsinized, harvested and swelled in 75 mM KCl for 10-15min at 37 °C. Subsequently, Cells were fixed with freshly made Carnoy’s solution (75% methanol, 25% acetic acid) on ice for 30 minutes. After washing with the fixative four times, cells were dropped onto glass slides and dried at room temperature. Slides were stained with DAPI washed briefly with PBS. For spreading HeLa cells expressing LAP-CENP-A and HeLa cells expressing RNaseH1/RNaseH1-2R, mitotic cells were obtained by mitotic shake-off and swelled in hypotonic buffer (75 mM KCl:0.8% NaCitrate:H2O at 1:1:1) with protease inhibitor cocktail (Roche) at room temperature for 10-15 min. Cells were spun to slides by Cytospin at 1500rpm for 5 min. The chromosome spreads were fixed with 2% PFA/PBS at room temperature for 20 min, then stained using the above protocol and indicated antibody concentrations. After washing with PBS, DNA was counterstained with DAPI.

**Table 1.** Antibody table.

| Epitope | Company | Catalogue # | Western Blot | Immunofluorescence | ChIP/DRIP |
| --- | --- | --- | --- | --- | --- |
| Flag | Sigma | F7425 | 1:10,000 | N/A | N/A |
| HA | Bethyl | A190-108 | 1:10,000 | N/A | N/A |
| Aurora B | Bethyl | A300-431A | 1:5,000 | 1:300 | N/A |
| Survivin | Cell Signaling | 2808 | 1:10,000 | N/A | N/A |
| RBMX | Cell Signaling | 14794S | 1:1,000 | 1:300 | N/A |
| Cyclin-B1 | Santa Cruz | Sc-594 | 1:1,000 | N/A | N/A |
| Sgo1 | Abcam | Ab58023 | 1:1,000 | 1:100 | N/A |
| Tubulin | ATCC Hybridoma | DM1a | 1:10,000 | 1:1,000 | N/A |
| H3 S10-phos | Millipore | 06-570 | 1:5,000 | 1:1,000 | N/A |
| SMC3 | Gift from S Rankin<br>(OK Medical research<br>Foundation) | N/A | 1:1,000 | N/A | N/A |
| Aurora B<br>(AIM1) | BD Biosciences | 611082 | N/A | 1:200 | N/A |
| Aurora B | Abcam | ab2254 | N/A | 1:200 | N/A |
| Anti-centromere<br>antigen (ACA) | Antibodies Inc | 15-234-0001 | N/A | 1:1,000 | N/A |
| H2ApT120 | Active Motif | 61195 | N/A | 1:1,000 | N/A |
| CENP-A | Abcam | ab13939-50<br>Clone3-19 | N/A | 1:1,000 | N/A |
| H3pT3 | Millipore | 07-424 | N/A | 1:500 | N/A |
| Aurora-BpT232 | Rockland Antibodies<br>and Assays | 600-401-677 | N/A | 1:200 | N/A |
| Borealin | Stukenberg lab | #968 | N/A | 1:200 | N/A |
| Bub1 | Abcam | ab54893 | N/A | 1:400 | N/A |
| R-loops | ATCC hybridoma | S9.6 | N/A | 1:1,000 | 1:10 |
| CENP-T | Foltz lab | N/A | N/A | 1:1,000 | N/A |
| mCherry | abcam | Ab183628 | 1:1000 | 1:200 | N/A |
| GFP | Stukenberg Lab | #786 | 1:1000 | N/A | 1:10 |
| GFP | abcam | 1218 | 1:1000 | 1:200 | N/A |

### ChIP

Chip analysis was performed as previously described. Briefly, cellular proteins and DNA were cross-linked by adding formaldehyde to the growth media to a final concentration of 0.1%. Cells were harvested in ice-cold phosphate-buffered saline and lysed with SDS buffer (50 mM Tris, 10 mM EDTA, and 1% w/v SDS). Lysates were sonicated utilizing a Branson sonifier 250 (Branson Ultrasonics, Danbury, CT) and precleared with salmon sperm DNA/protein A-agarose (Upstate Biotechnologies, Lake Placid, NY). Lysates were then tumbled overnight at 4 °C with salmon sperm DNA/protein A-agarose with anti-GFP or rabbit IgG antibodies. Complexes were precipitated and serially washed three times each with low salt (20 mM Tris, 150 mM NaCl, 2 mM EDTA, 0.1% (w/v) SDS, and 1% (v/v) Triton X-100); high salt (20 mM Tris, 500mM NaCl, 2 mM EDTA, 01% (w/v) SDS, and 1% (v/v) Triton X-100); LiCl wash (10 mM Tris, 250 mM LiCl, 1 mM EDTA,1%(w/v) deoxycholate, and 1% (v/v) Nonidet P-40); and TE buffer (20 mM Tris and 2 mM EDTA). Washed complexes were eluted with freshly prepared elution buffer (1% SDS and 100 mM NaHCO3), and the Na+ concentration was adjusted to 200 mM by adding NaCl followed by incubation at 37 °C to reverse protein/DNA cross-links. DNA was purified utilizing a PCR purification kit (Qiagen). Purified DNA was then amplified across the il8 locus region or centromeric α-satellite DNA on chromosome 7, primers available in Table 3.

### Proximity ligation assay (PLA)

PLA was performed as described (Banerjee et al., 2014).

### DRIP/DRIP-seq

DLD1 cells were arrested in colcemid for 24 hours and then treated with either DMSO, AZD-1152 or ZM-447439 at the indicated concentrations (Table 2) for 1 hour. Mitotic shakeoffs were performed to gain a mitotic population. DRIP assays were performed as in Halász et al., 2017. Briefly, cells were fixed using 1% Formaldehyde for 10 minutes, then quenched with Glycine to a final concentration of 0.5 M at room temperature. Cells were collected, washed twice with PBS, resuspended in lysis buffer (50 mM HEPES-KOH, pH 7.5, 140 mM NaCl, 1 mM EDTA, 1% Triton X-100, 0.1% Na-Deoxycholate, 1% SDS), 1 mL per 10^7^ cells, lysed by being passed through a 20G needle 10 times, and sonicated 15 cycles of 30s on, 30s off, High setting, Bioruptor. This yielded an average of 300 bp fragment. Sonicated chromatin was digested with Proteinase K to 1 μg/ml at 65 °C overnight to remove proteins and crosslinks. DNA was precipitated using 1/10 volume 3 M Na-acetate and 1 volume isopropanol and incubated for 1 hour at −80 °C. The DNA pellet was washed, dried, and resuspended in 100 μl 5 mM Tris-HCl pH 8.5. 12 μg of the resulting DNA was incubated with RNaseH1 buffer +/- RNaseH1 (40 units, Takara) overnight at 37 °C, then 10 μg of the reaction was incubated with 5 μg of S9.6 hybridoma antibody overnight in IP buffer (50 mM HEPES-KOH pH 7.5, 140 mM NaCl, 5 mM EDTA, 1% Triton X-100, 0.1% Na-Deoxycholate), rotating at 4 °C. 25 μl pre-blocked Dynabeads Protein A (Thermo Fisher, blocked for 1 hour in PBS/EDTA with 0.5% BSA) were added to the immunoprecipitation and rotated for 4 hours at 4 °C. Beads were recovered and washed successively for 30 mins at room temperature each: 2 washes of 1 ml low salt buffer (50 mM HEPES-KOH pH 7.5, 140 mM NaCl, 5 mM EDTA, 1% Triton X-100, 0.1% Na-Deoxycholate), 2 washes of 1 ml high salt buffer (50 mM HEPES-KOH pH 7.5, 500 mM NaCl, 5 mM EDTA, 1% Triton X-100, 0.1% Na-Deoxycholate), 2 washes of 1 ml LiCl wash buffer (10 mM Tris-HCl pH 8, 250 mM LiCl, 1 mM EDTA, 1% Triton X-100, 0.1% Na-Deoxycholate, 0.5% NP-40), and 2 washes of 1 ml TE buffer (10 mM Tris-HCl pH 8, 10 mM EDTA). Elution was performed in 100 μl elution buffer (50 mM Tris-HCl pH 8, 10 mM EDTA, 1% SDS) for 15 minutes at 65 °C, vortexing every 5 minutes. Resulting supernatant DNA was purified using a PCR clean-up kit (Invitrogen), along with 1 μg starting DNA from the RNaseH1 reaction. The recovered DNA was analyzed by quantitative real-time PCR using LunaScript qPCR mastermix (NEB) and the ABI StepOnePlus qPCR machine and primers in Table 3. 45 ng of precipitated DNA spiked with 5 ng of fragmented genomic DNA from *S. cerevisiae* (strain MT11, a gift from the David Auble) was used as starting material for the Takara SMARTer ThruPLEX DNA-seq Kit and DNA HT Dual Index Kit. Libraries were sequenced using the Illumina NextSeq 500 at the UVA Genome Analysis and Technology Core using 12-plex multiplexing, mid output 150 round paired end sequencing, resulting in greater than 10M reads per sample.

**Table 2.** Chemical and protein inhibitors

| Inhibitor | Company | Catalogue # | Concentration<br>HeLa cells | Concentration<br>RPE1 cells | Concentration<br>DLD1 cells |
| --- | --- | --- | --- | --- | --- |
| ZM447439 | Selleck Chem | S1103 | 2 μM | 4 μM | 2 μM |
| AZD1152 | Cayman Chemical | 13647 | 500 nM | 1 μM | 500 nM |
| Hesperadin | Selleck Chem | S1529 | 100-500 nM | N/A | N/A |
| Colcemid | Gibco | 15212012 | 100 ng/ml | 200 ng/ml | 100 ng/ml |
| Nocodazole | Sigma | M1404 | 0.33-3.3 μM | N/A | N/A |
| Thymidine | Sigma | T1895 | 2 mM | 4 mM | N/A |
| RNaseH1 | Takara | 2150B | N/A | N/A | 40 U/ 10 μg DNA |

**Table 3.**
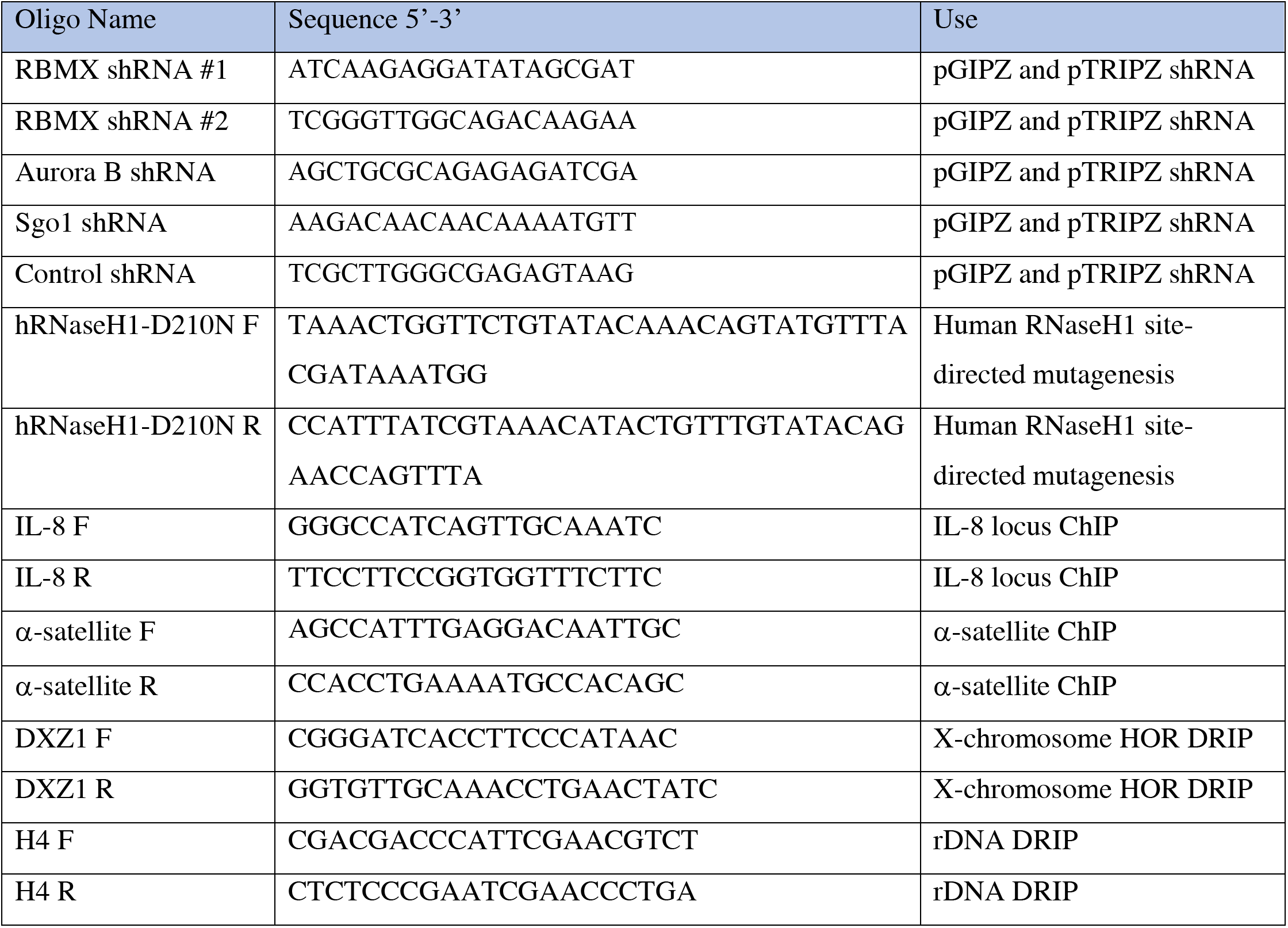
Oligonucleotides.

### Bioinformatic Analysis

HEK293 reads were obtained from the European Nucleotide Archive (ENA) under accession number GSE68953 for the study (Nadel et al., 2015). These reads were processed identically to the reads derived in this study. Reads were quality thresholded using PRINSEQ (Minimum mean quality = 15, minimum quality to trim = 20, minimum length to drop = 100, maximal N percentage = 2) and subsampled to 6.5M reads using seqtk. Sequences were first aligned to SacCer3 genome using BWA-MEM (Li and Durbin, 2009) to determine spike percentage (HEK293 reads used as background alignment for sequence conservation) in order to determine a scaling factor according to library preparation efficiency. These values are listed below in Table 4. Due to an error in de-multiplexing, R2 file of ZM sample had to be reverse complemented to generate FR oriented pairs. Reads were aligned to the human genome (hg38) using BWA-MEM. Given that we expected to observe repetitive sequences in our samples, we did not use repeat-masking but required only one alignment per read pair. Peaks were called using MACS2 (Feng et al., 2012; Zhang et al., 2008), with replicate samples used to refine peaks common to both replicates (FDR 1%, ZM and AZD were considered replicates in this case). Tag Directories were created using Homer makeTagDirectory (Heinz et al., 2010) and were used to generate enrichment graphs using annotatePeak (Heinz et al., 2010) using the peak profile setting. Homer peak annotation was used to generate genome ontology graphs. The most recent GENCODE version (v32, Frankish et al., 2019) was used for annotation of gene bodies, transcription start sites (TSS) and transcription termination sites (TTS). RepeatExplorer Galaxy instance (Novák et al., 2013) was used to generate *de novo* repeat clusters using compiled input files. The subsampled files were then aligned to the contigs generated using a BLAST-N similarity search (Neumann et al., 2012) to gain a count table, which was then normalized, scaled and repeat experiments were averaged. HOR specific K-mers were obtained from the supplemental information in Miga, 2017. Kmer counts were obtained using the KAT sect function (Mapleson et al., 2017).

**Table 4.** Spike in quantification.

| Sample | Reads Mapped and Paired | Corrected | Scaling factor |
| --- | --- | --- | --- |
| Input 1 | 2.94M | 2.84M | 0.211 |
| Input 2 | 2.15M | 2.05M | 0.293 |
| HEK293 Input | 0.10M | 0 | 0.9* |
| DMSO 1 | 3.75M | 3.48M | 0.172 |
| DMSO 2 | 1.32M | 1.05M | 0.571 |
| AZD | 2.07M | 1.80M | 0.333 |
| ZM | 1.25M | 0.98M | 0.612 |
| DMSO-RNH1 1 | 4.63M | 4.36M | 0.138 |
| DMSO-RNH1 2 | 5.52M | 5.25M | 0.114 |
| HEK293 DRIP | 0.27M | 0 | 0.9* |

\*Assumed complete efficiency with library preparation

### Purification of mitotic chromatin and MNase digestion

Either LAP, LAP-Aurora B, LAP-Survivin or LAP-Borealin expressing HeLa cells (3×109) were arrested with colcemid for 16-18 hours were harvested and mitotic chromosomes were purified as described (Paulson, 1982). Mitotic chromosomes were resuspended in 40ml MNase buffer (20mM HEPES, pH7.7; 20mM KCl; 5mM β-mercaptoethanol; 1xProtease Inhibitors cocktail (EDTA free, Roche), 20mM β-glycerophosphate,1mM Sodium orthovanadate, 3mM CaCl2, 250 mM NaCl, and 0.1% Digitonin) and digested with MNase (Roche, 150u ml-1) for 1 hour at room temperature. After digestion extracts were supplemented with 50 mM NaF and clarified by centrifugation at 12,000g for 20 minutes at 4°C. The supernatants were used as starting materials for LAP purifications that are described in the Figure 1–figure supplement 1.

### Multidimensional Protein Identification Technology (MudPIT)

The elutes from LAP purifications were digested in solution using trypsin. The digested samples were pressure-loaded onto a fused silica capillary desalting column containing 5 cm of 5 μm Polaris C18-A material (Metachem, Ventura, CA) packed into a 250-μm i.d. capillary with a 2 μm filtered union (UpChurch Scientific, Oak Harbor, WA). The desalting column was washed with buffer containing 95% water, 5% acetonitrile, and 0.1% formic acid. After desalting, a 100 μm i.d capillary with a 5-μm pulled tip packed with 10 cm 3 μm Aqua C18 material (Phenomenex, Ventura, CA) followed by 3 cm 5-μm Partisphere strong cation exchanger (Whatman, Clifton, NJ) was attached to the filter union and the entire split-column (desalting column–filter union–analytical column) was placed in line with an Agilent 1100 quaternary HPLC (Palo Alto, CA) and analyzed using a modified 12-step separation described previously (Washburn et al., 2001). As peptides eluted from the microcapillary column, they were electrosprayed directly into an LTQ 2-dimensional ion trap mass spectrometer (ThermoFinnigan, Palo Alto, CA) with the application of a distal 2.4 kV spray voltage. A cycle of one full-scan mass spectrum (400-1400 m/z) followed by 8 data-dependent MS/MS spectra at a 35% normalized collision energy was repeated continuously throughout each step of the multidimensional separation. Application of mass spectrometer scan functions and HPLC solvent gradients were controlled by the Xcalibur datasystem. MS/MS spectra were analyzed using the following software analysis protocol. Poor quality spectra were removed from the dataset using an automated spectral quality assessment algorithm (Bern et al., 2004). MS/MS spectra remaining after filtering were searched with the SEQUEST™ algorithm (Eng et al., 1994) against the current version of NCBI Homo sapiens database concatenated to a decoy database in which the sequence for each entry in the original database was reversed (Peng et al., 2003). SEQUEST results were assembled and filtered using the DTASelect (version 2.0) program.

### Statistical Tests

All statistical tests were run with the assistance of the Graphpad Prism software. First, statistical outliers were determined and mathematically eliminated using the ROUT method, with Q value of 1%. For all measurements, descriptive statistics were then used to determine whether the data conformed to a normal distribution. For the data in which all samples passed a D’Agostino and Pearson test (K2 value measured), data were considered to be normally distributed and a parametric test was applied. These parametric tests were either a student’s t-test for comparisons between two samples with similar standard deviations, or Welch’s t-test for comparisons between samples with at least 2-fold differing standard deviations, or a one-way ANOVA for comparison of multiple samples. In the case of ANOVA, Dunnet correction for multiple comparison was utilized, and a q statistic was measured for each difference. If at least one sample in a dataset involved non-normally distributed data, non-parametric tests were applied. For comparison between two samples, Mann-Whitney tests were used to compare the ranks of individual data points within the total distribution, a Mann-Whitney U value was collected, and a two-tailed p-value was determined. For comparison of multiple unpaired samples, a Kruskal-Wallis test was performed to compare ranks of individual data points within the total distribution. Dunn’s correction for multiple comparisons was performed on post-test statistics, and Z statistics were used to determine approximate p-values. Two-sided p-values were always determined unless stated within figure legends. If measurements could not be taken for each comparison directly, multiple comparison corrections were applied. Figures show means and ranges of data if normally distributed, medians and ranges of data if not normally distributed.

---

**Supplemental Figure 1.**
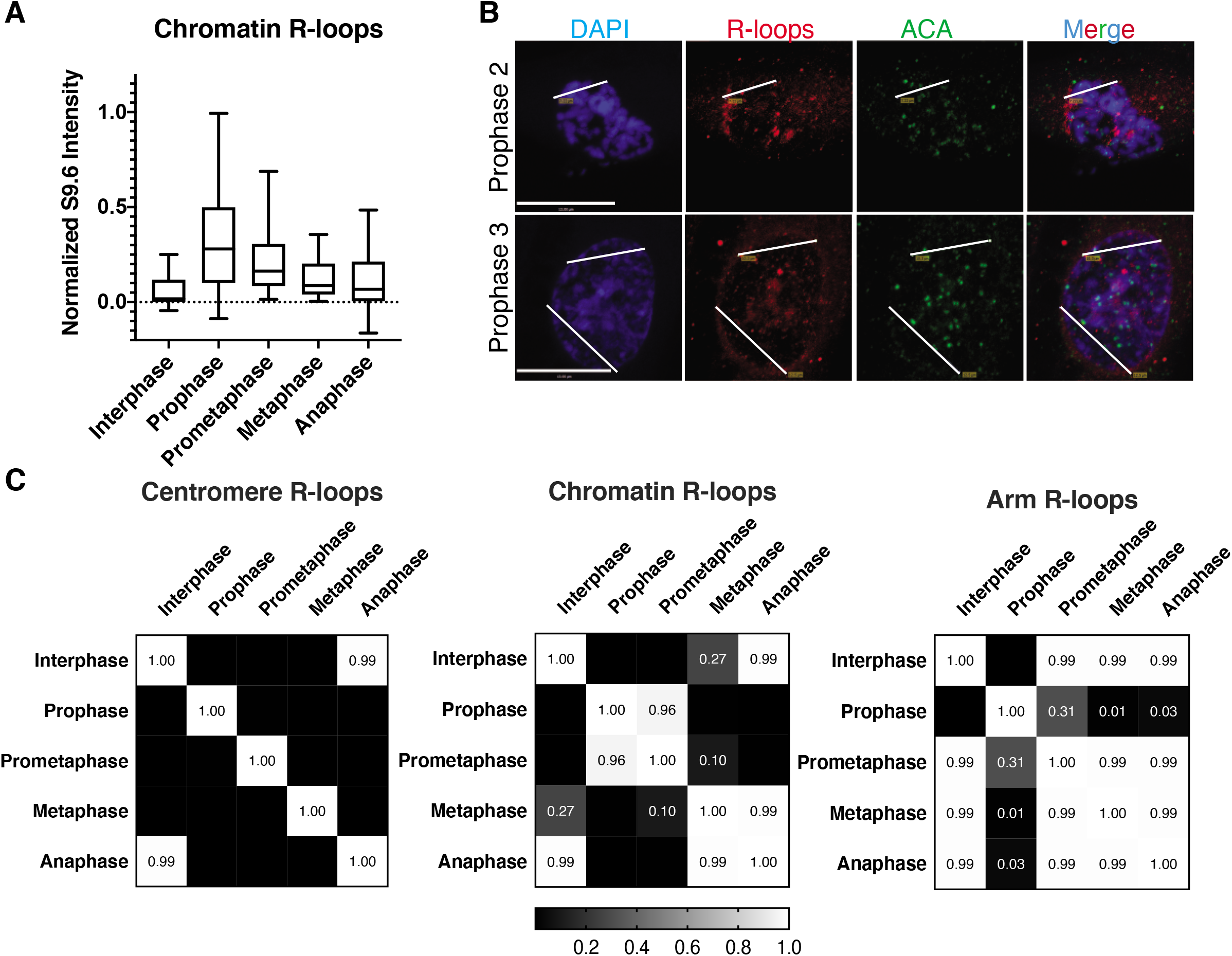
R-loops are associated with condensing chromatin. A. Total chromatin intensities of R-loops, quantified by total S9.6 signal over the DAPI stained region, normalized to DAPI. B. Additional images of prophase nuclei with additional line scan tracks from figure 1E demarcated. Scale bars, 13 μm. C. Significance values of S9.6 quantifications from Kruskal-Wallis non-parametric ANOVAs with post-test to estimate approximate p-values, given that data in all cases failed multiple normality tests. Black boxes, p<0.0001; all other values listed within the heatmaps.

**Supplemental Figure 2.**
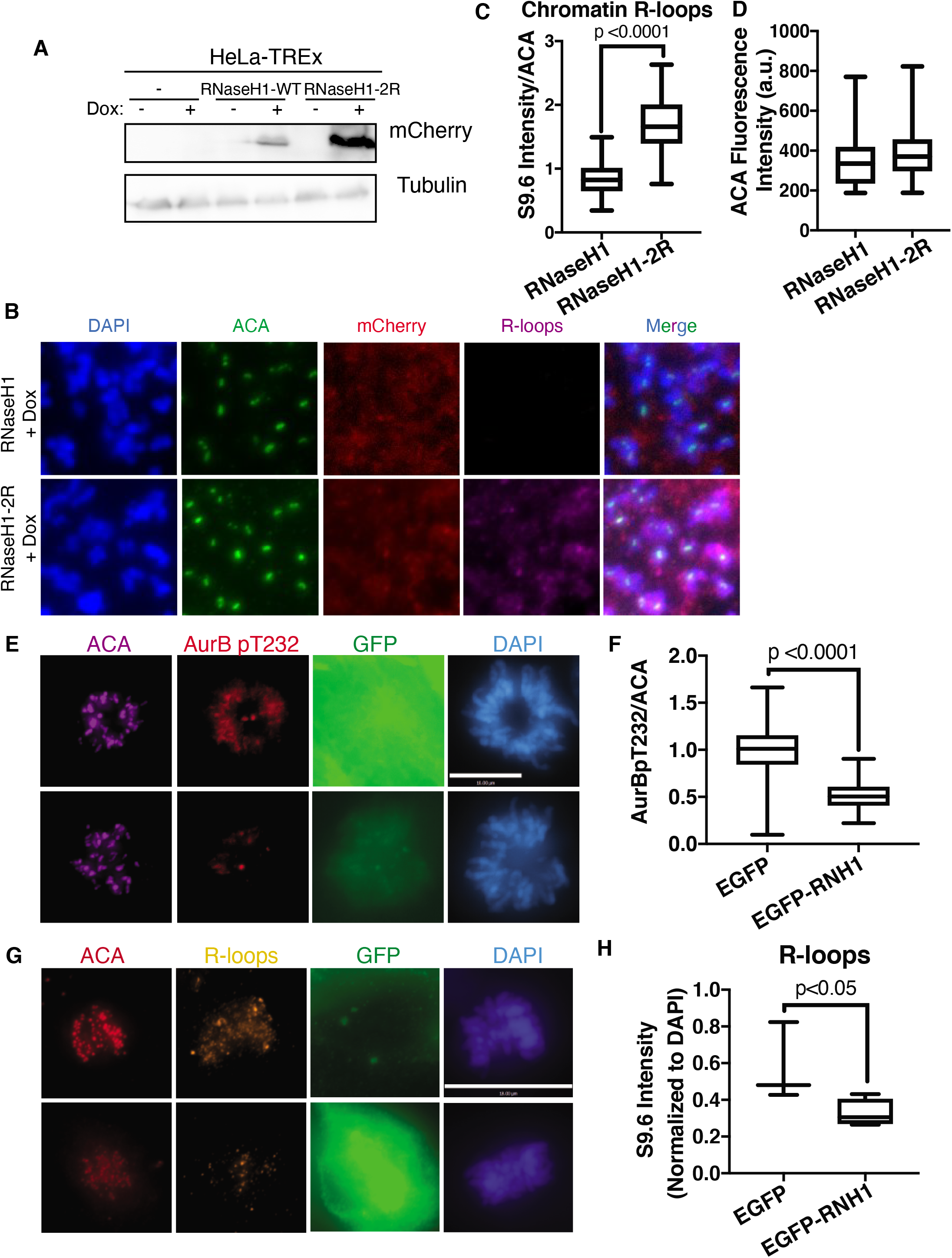
RNaseH1 overexpression controls R-loop prevalence, Aurora B activation. A. Western blot showing induction of mCherry-RNaseH1 constructs after 8 hours of doxycycline addition. B. Mitotic spreads of HeLa-T-REx cells overexpressing mCherry-RNaseH1 constructs, with indirect immunofluorescence for ACA, mCherry, and S9.6. C. Quantification of S9.6 normalized to ACA. D. Quantification of total ACA signal, not significantly different in the two conditions. E-H. RPE1-T-REx cells treated as in Figure 1 E-H, with indirect immunofluorescence for ACA and Aurora B pT232 (E-F) or ACA, S9.6, and GFP (G-H). F. Quantification of Aurora B pT232 normalized to ACA. H. Quantification of S9.6 normalized to ACA.

**Supplemental Figure 3.**
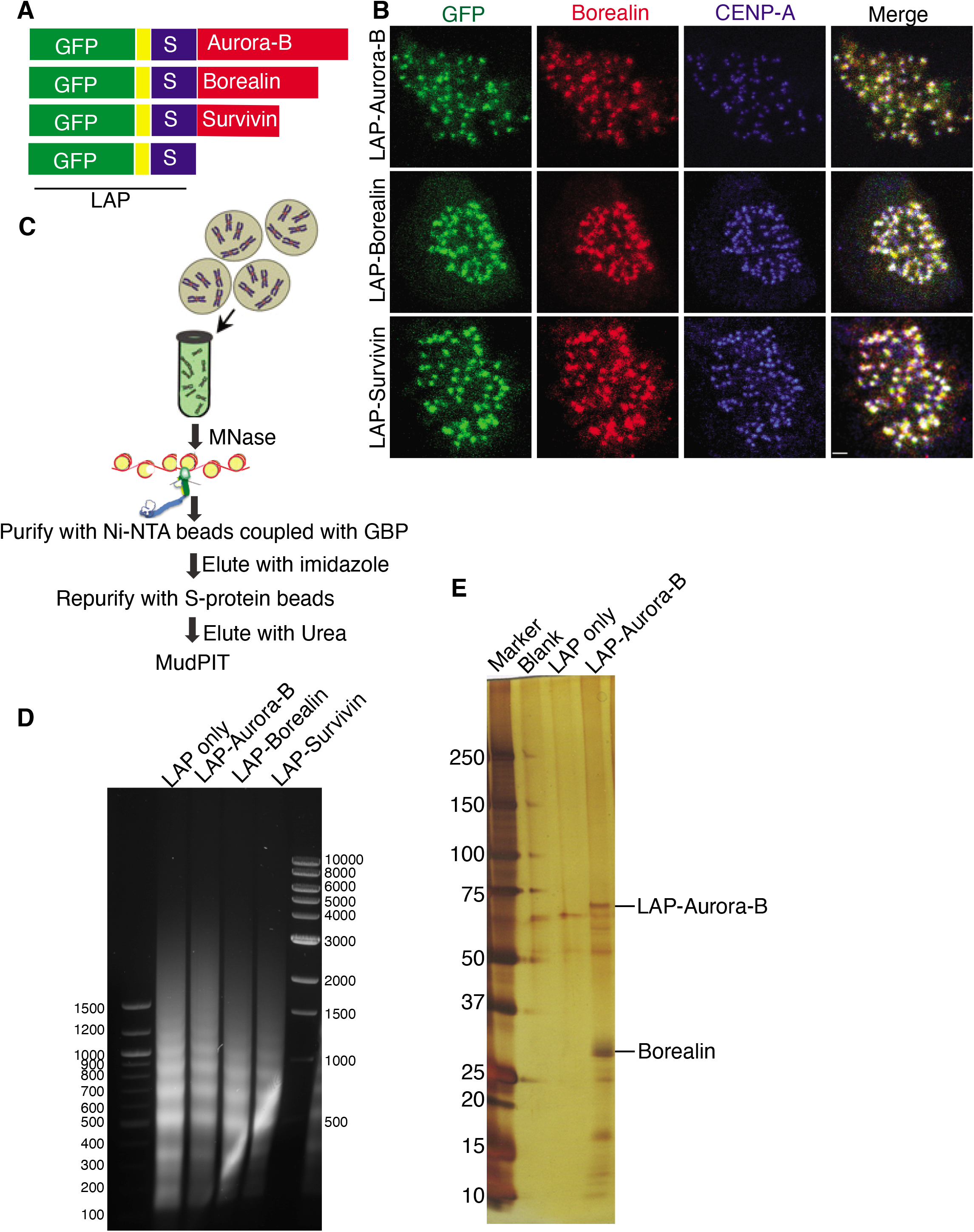
Validation of MudPiT analysis of CPC members. A. Schematic representation of constructs expressed in HeLa-T-REx cells. B. Validation of localization of the LAP-tagged proteins. Overlap with indirect immunofluorescence of Borealin in mitotic cells indicates that the proteins localize correctly. C. Schematic representation of purification of proteins. D. DNA fragmentation after Micrococcal Nuclease (MNase) treatment. E. Silver stain gel showing purification of CPC members from LAP-Aurora B purification.

**Supplemental Figure 4.**
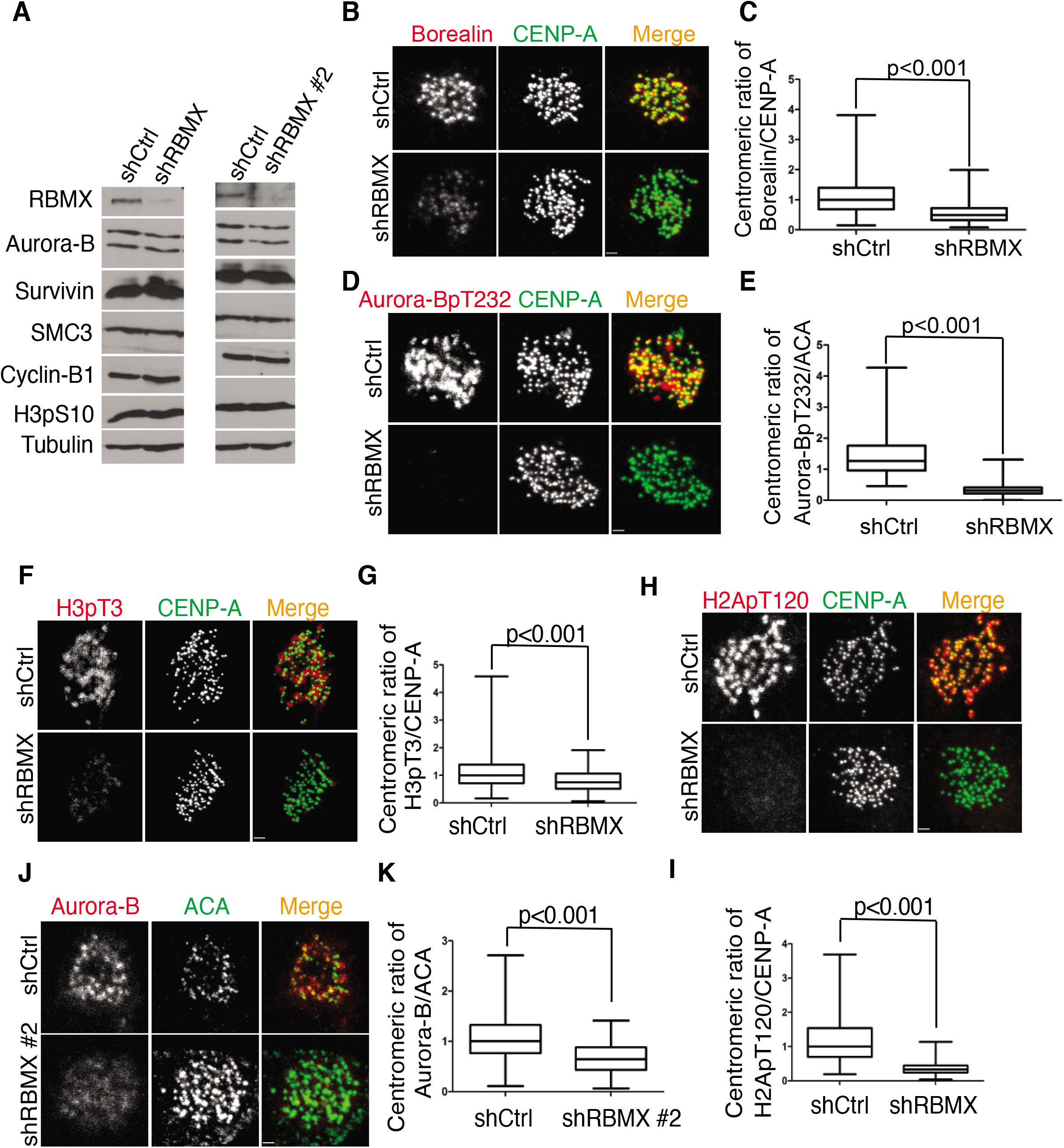
RBMX is necessary to localize the CPC and CPC localization signals. A. Two different shRNA constructs against RBMX effectively knock down protein levels of RBMX but have no effect on protein levels of CPC members, cohesion, Cyclin B1, or H3S10 phosphorylation. B-I. Knockdown of RBMX by the first shRNA in A. in HeLa-T-REx cells leads to loss of B. Borealin immunofluorescence, quantified in C., D. Aurora B pT232 immunofluorescence, quantified in E., F. H3pT3 immunofluorescence, quantified in G., and H. H2ApT120 immunofluorescence, quantified in I. J, knockdown of RBMX by the second shRNA confirms loss of Aurora B immunofluorescence, quantified in K. P-values determined by unpaired two-tailed t-test.

**Supplemental Figure 5.**
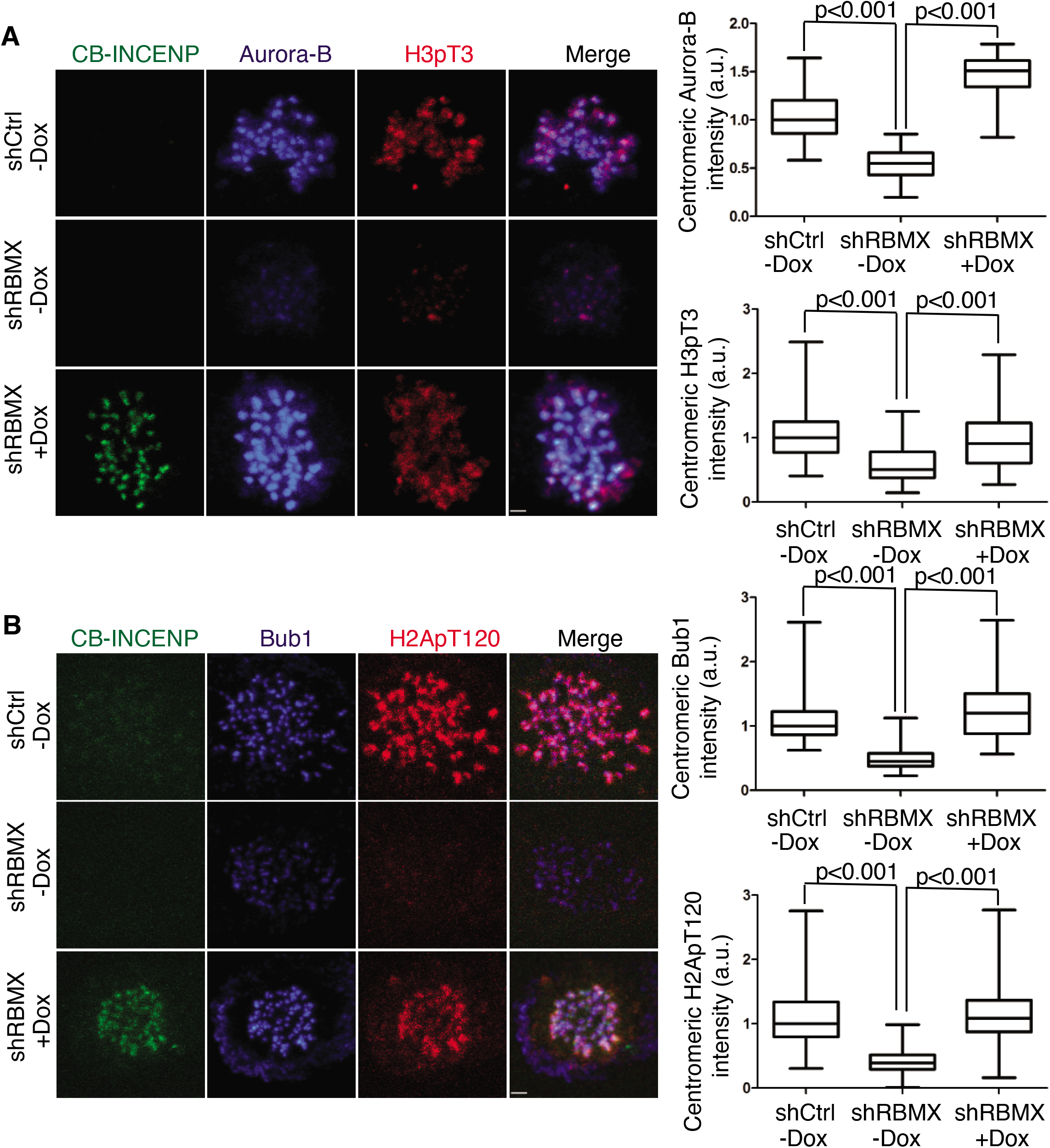
RBMX is necessary to localize the CPC and CPC localization signals in an Aurora B activity dependent manner. A. Representative images of cells knocking down RBMX using shRNA #1 and adding back CENP-B DNA binding domain fused to INCENP in a tet-inducible manner, stained using indirect immunofluorescence for Aurora B and H3pT3. Centromeric Aurora B quantified in top left, centromeric H3pT3 quantified in bottom left. B. Representative images of cells knocking down RBMX using shRNA #1 and adding back CENP-B DNA binding domain fused to INCENP in a tet-inducible manner, stained using indirect immunofluorescence for Bub1 and H2ApT120. Centromeric Bub1 quantified in top left, centromeric H2ApT120 quantified in bottom left.

**Supplemental Figure 6.**
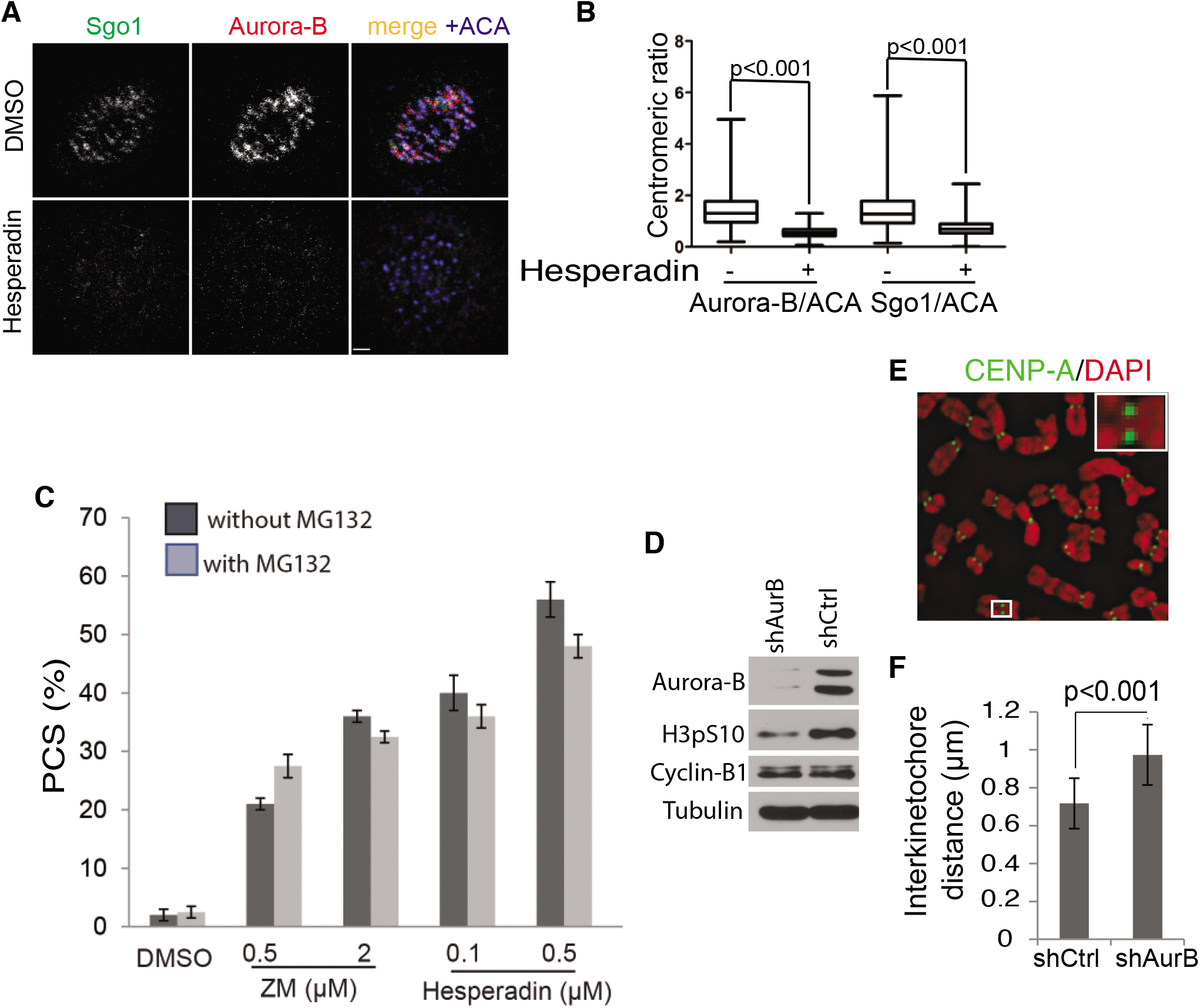
Aurora B controls localization of Sgo1, centromeric cohesion. A. HeLa-T-REx cells treated with Aurora B inhibitor Hesperadin at 100 nM and stained using indirect immunofluorescence for Aurora B, Sgo1 and ACA. Fluorescence intensity normalized to ACA is quantified in B, p-values determined by unpaired two-tailed t-test. C. PCS assay utilizing two different Aurora B inhibitors, ZM-447439 or Hesperadin, with and without MG132. D. Western blots validating knockdown of Aurora B using an shRNA against Aurora B, relative to non-targeting shCtrl. E. Interkinetochore distance assay, utilizing HeLa-T-REx cells stably expressing LAP-CENP-A either with shCtrl or shAurora B. Increase in interkinetochore distance quantified in F, p-value determined by unpaired two-tailed t-test.

## Acknowledgements

We would like to acknowledge Dr. Sathyan Kizhake Mattada, Dr. Michael Guertin, Dr. Manikarna Dinda, and Dr. David Auble for reagents, Dr. Dan Burke for helpful comments and manuscript review, and Dr. Katherine Pfister, Dr. Ewa Niedzialkowska, Dr. Pawel Janczyk, Tan Truong, and Luke Elderidge for helpful discussions. Funding was provided by the National Institutes of Health grant P41 GM103533 (JY), T32 CA 9109-43 (EM), R01 GM118798 (PTS) and R01 GM124042 (PTS).

